# Massively Parallel Free Energy Calculations for *in silico* Affinity Maturation of Designed Miniproteins

**DOI:** 10.1101/2024.05.17.594758

**Authors:** Dylan Novack, Si Zhang, Vincent A. Voelz

**Affiliations:** Department of Chemistry, Temple University, Street, Philadelphia, 19122, PA, USA

**Author notes:** Contributing authors.

**Keywords:** computational protein design, site-saturation mutagenesis, affinity maturation, free energy methods, Rosetta, *de novo* design, hemagglutinin, HA, influenza, expanded ensemble, molecular simulation

## Abstract

Computational protein design efforts continue to make remarkable advances, yet the discovery of high-affinity binders typically requires large-scale experimental screening of site-saturated mutant (SSM) libraries. Here, we explore how massively parallel free energy methods can be used for *in silico* affinity maturation of *de novo* designed binding proteins. Using an expanded ensemble (EE) approach, we perform exhaustive relative binding free energy calculations for SSM variants of three miniproteins designed to bind influenza A H1 hemagglutinin by Chevalier et al. (2017). We compare our predictions to experimental **ΔΔ*G*** values inferred from a Bayesian analysis of the high-throughput sequencing data, and to state-of-the-art predictions made using the Flex ddG Rosetta protocol. A systematic comparison reveals prediction accuracies around 2 kcal/mol, and identifies net charge changes, large numbers of alchemical atoms, and slow side chain conformational dynamics as key contributors to the uncertainty of the EE predictions. Flex ddG predictions are more accurate on average, but highly conservative. In contrast, EE predictions can better classify stabilizing and destabilizing mutations. We also explored the ability of SSM scans to rationalize known affinity-matured variants containing multiple mutations, which are non-additive due to epistatic effects. Simple electrostatic models fail to explain non-additivity, but observed mutations are found at positions with higher Shannon entropies. Overall, this work suggests that simulation-based free energy methods can provide predictive information for *in silico* affinity maturation of designed miniproteins, with many feasible improvements to the efficiency and accuracy within reach.

## 1 Introduction

*De novo* protein design is a powerful tool that promises transformative advances in biotechnology from biomedical therapeutics to nanomaterials.[1] Using computational algorithms alongside high-throughput experiments, it is now possible to design mini-protein binders to arbitrary targets.[2, 3] While impressive, the success rate remains low: many potential designs must be screened experimentally to discover low-affinity binders. Identifcation of high-affinity binders typically requires screening site-saturation mutagenesis (SSM) libraries of the low-affinity designs.[3, 4] This indicates that the energy function and/or conformation sampling is apparently not accurate enough to be used for *in silico* affinity maturation prediction, a limitation that is likely due to the low-resolution models used for electrostatics and solvation in tools like Rosetta.

Meanwhile, alchemical free energy approaches utilizing all-atom simulations have shown remarkable success in predicting relative binding affinities.[5–9]Recent benchmarks from free energy perturbation (FEP),[10, 11] non-equilibrium work,[12] and multisite *λ* dynamics[13] approaches suggest that relative binding free energies (RBFE) for single-residue mutations in protein-protein interfaces can be predicted within an accuracy of 2 kcal/mol. Can all-atom free energy approaches more accurately predict SSMs? If so, it would be an important step toward a completely computational protocol to generate *de novo* binders.

To explore this idea, we perform a benchmark study to compare *in silico* SSM predictions from alchemical free energy calculations with experimental affinity maturation studies from Chevalier et al., in which massively parallel *de novo* design using Rosetta was used to discover high-affinity protein binders of the highly conserved stem region of influenza A H1 hemagglutinin (HA).[4] Tens of thousands of binders derived from mini-protein scaffolds were computationally designed and experimentally screened using yeast display, fluorescence activated cell sorting (FACS), and next-generation sequencing. The best binders were then used as starting points to generate SSM libraries for similar high-throughput screening. Finally, the SSM results were then used to guide the further generation of a diverse library of binders incorporating favorable mutations and error-prone PCR, to identify affinity-matured mini-proteins that bind in the low nanomolar range.

While Chevalier et al. published SSM results for several miniprotein binders (of HA and botulinum neurotoxin B binders), here we focus on three noteworthy designs: A8, A13, and A18 (Figure 1). These 40-residue miniproteins all have a EHEE topology, and bind HA with an apparent *K*_*d*_ over 300 nM. Affinity-matured variants of these designs each contain 5, 6, or 7 mutations that confer a significant increase in affinity (*K*_*d*_ ≤6 nM). In each case, the majority of the mutations occur away from the binding interface, raising questions about how such affinity is achieved. The full SSM results entail (19 amino acids ×[38 + 38 + 37] positions for the three designs) = 2147 single-point mutations.

**Fig. 1:**
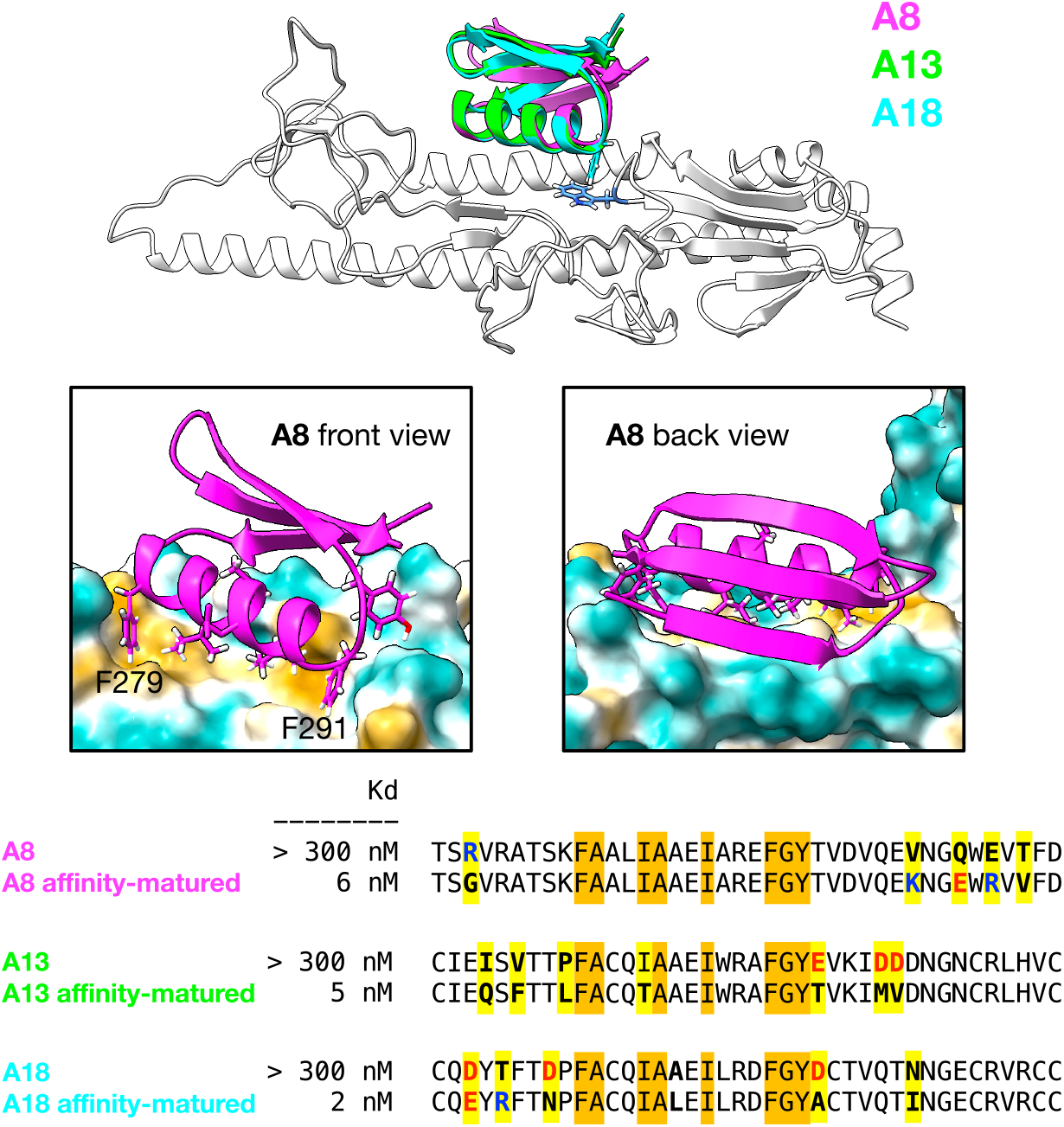
Computationally designed miniproteins that bind the hemagglutinin (HA) stem region. (top) Structural models from Chevalier et al.[4] of miniproteins A8 (magenta), A13 (green) and A18 (cyan) bound to a truncated model of HA (white) constructed from PDB:3R2X. Shown in blue is Trp119 from HA, which interacts with F291 (shown for A18) on each miniprotein, anchoring the binding motif. (middle) Front and back views of A8 modeled in complex with HA, shown with a hydrophobic surface showing nonpolar (orange) and polar (teal) regions. Residues F279 and F291 flank the conserved binding helix. (bottom) Experimental affinities and sequences of miniproteins and their affinity-matured variants. Highlighted in orange are conserved residues across all variants that comprise the conserved binding helix motif. Highlighted in yellow are positions that are mutated upon affinity-maturation, colored by net charge. ChimeraX was used for molecular visualization.

### Expanded ensemble methods enable massively parallel free energy calculations

To efficiently perform *in silico* SSM screens using modern alchemical free energy calculations, we take advantage the method of expanded ensembles (EE),[14–16] which allows relative free energies to be calculated on a massively parallel scale. While many alchemical FEP methods require multiple tightly coupled simulation replicas, EE simulations are able to sample all alchemical intermediates in a single simulation, making it ideal for asynchronous client-server distributed computing via the Folding@home platform.

Alchemical EE approaches have been underutilized in the past because of convergence problems that arise from non-optimal choices of alchemical intermediates and by slow conformational dynamics. The first problem has largely been addressed (see Methods) while the second problem is an area of active research.[17, 18] Hence, large-scale *in silico* SSM screens provide an excellent evaluation of current EE approaches for this purpose.

In this work, we use EE methods to perform massively parallel *in silico* SSM screens for mini-protein HA binders A8, A13 and A18. We characterize the convergence and uncertainty estimates of these predictions, and compare their accuracy against experiment. Since Chevalier et al. only report binding enrichment fractions from sequence reads, we use Bayesian inference to infer apparent *K*_*d*_ values of all binders, allowing for a more quantitative comparison of relative binding free energies. We also compare to state-of-the-art estimates of relative binding free energies from Rosetta using the Flex ddG algorithm.[19]

In interpreting these comparisons, it is important to keep in mind that the structures of A8, A13 and A18 we use are themselves computational models predicted by Rosetta. Our comparisons test the larger question of whether alchemical free energy estimates can produce useful information to guide the selection of higher-affinity designs in the absence of experimental structural information. These additional layers of uncertainty suitably reflect the practical challenges involved in computational protein design.

Finally, we use the inferred apparent *K*_*d*_ values from the experimental SSM studies to help understand the set of mutations that lead to affinity-matured versions of A8, A13 and A18.

## 2 Results and Discussion

### 2.1 Expanded ensemble estimates of ΔΔ*G* for site-saturated mutants of *de novo* designed hemagglutinin binders

Expanded ensemble (EE) calculations were performed using GROMACS 2020.3[20] on Folding@home.[21] Starting from the predicted structural models of A8, A13 and A18 from Rosetta,[4] alchemical hybrid topologies were constructed for all possible single-residue mutations using *pmx*.[22] The AMBER14sb[23] force field was used for the protein along with the TIP3P water model. Backbone heavy atom restraints were used for both HA and miniprotein.

Accurate estimates of relative binding free energies for mutations with net charge differences are a particular challenge.[24] We deal with this by adding one or more neutralizing counterions to the alchemical topology, restraining their positions in solution away from the protein(s).

In our EE approach,[15, 16, 25] a series of *N* = 45 alchemical intermediates were constructed, parameterized by *λ*_*i*_ ∈ [0, 1], for *i* = 1, …*N*. The *λ* value interpolates the non-bonded interactions between a *λ* = 0 topology for the wild-type residue and a *λ* = 1 topology for the mutant residues (Figure 2). A soft-core potential is used to avoid discontinuities.[26]

**Fig. 2:**
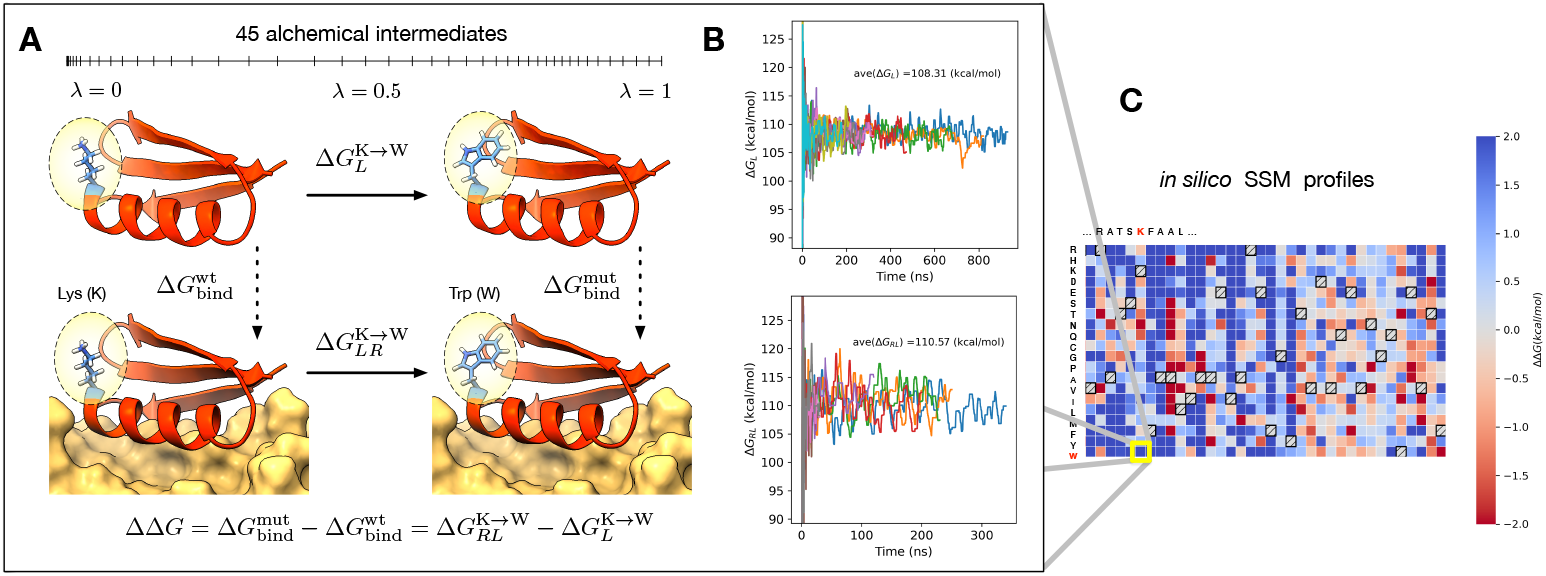
Overview of the *in silico* Site-Saturated Mutagenesis (SSM) protocol. (A) In this example, expanded ensemble simulations are used to estimate the free energy of transformation for K278W in solution (Δ*G*_*L*_) and in complex with HA (Δ*G*_*RL*_). Hash marks on the *λ*-axis show a typical optimized schedule of values. The relative free energy of binding is computed as ΔΔ*G* = Δ*G*_*RL*_− Δ*G*_*L*_. (B) To estimate each free energy, the mean result over ten replicates is used, where values are collected after a convergence criteria is reached. (C) Calculations are performed in parallel for all amino acids at all positions, to construct a complete SSM profile.

Throughout an EE simulation, Markov chain Monte Carlo (MCMC) transitions are proposed from thermodynamic ensemble *i* → *j*, and accepted according to a Metropolis criterion. A bias energy −*f*_*i*_ is applied to each ensemble, and adjusted adaptively throughout the simulation using the Wang-Landau flat-histogram algorithm[27] to equalize the acceptance probabilities in each direction. By detailed balance, the values 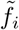 that achieve uniform sampling are the free energy of each thermodynamic ensemble *i*; therefore, the free energy of the *λ* = 0→1 transformation is given by 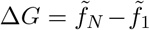. Critical to avoiding sampling bottlenecks across thermodynamic ensembles is the choice of the *λ*_*i*_ values. Poor choices of *λ*_*i*_ values will result in weak thermodynamic overlap between ensembles, and low MCMC acceptance probabilities, leading to slow convergence. To eliminate these bottlenecks, we first perform a pre-production EE simulation constrained to only nearest-neighbor thermodynamic transitions; the resulting distributions of (reduced) energy perturbations Δ*u*_*k,k*+1_ and Δ*u*_*k*+1,*k*_ are then used by *pylambdaopt* algorithm to minimize the thermodynamic length between adjacent ensembles (see Methods).

In an EE simulation, *N* thermodynamic ensembles corresponding to alchemical intermediates are adaptively sampled through proposed Markov Chain Monte Carlo (MCMC) moves. A bias potential 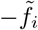 applied each intermediate *i* is maintained and adaptively adjusted using the Wang-Landau flat histogram method[27], with the goal of achieving uniform sampling of all intermediates. By detailed balance, the converged values of 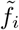 that achieve this goal are equal to the true free energies *f*_*i*_ of each thermodynamic ensemble, and the total free energy change along the alchemical transformation from *i* = 1 to *i* = *N* can be computed as Δ*G* = *f*_*N*_ −*f*_1_.

To estimate changes in the the free energy of binding ΔΔ*G* due to mutation, the free energy of two alchemical transformations need to be computed: Δ*G*_*L*_, the ligand-only mutational change for the mini-protein in solution, and Δ*G*_*RL*_, the receptor-ligand mutational change for the mini-protein bound to HA, where ΔΔ*G* = Δ*G*_*RL*_ − Δ*G*_*L*_ (Figure 2).

In practice, EE free energy estimates fluctuate over time due to the time-correlation inherent in conformational sampling, and the learning rate of the flat-histogram method. We have found, however, that improved estimates can be made by averaging estimates over multiple independent simulations, and only collecting data for these estimates until after certain convergence criteria are met.[15, 16] For this study, we perform 10 replicates of each EE simulation, and deem convergence to be reached when the Wang-Landau histogram increment is below 0.01 *k*_*B*_*T* ; if none of the trajectories satisfy this criterion, we use a secondary criterion of 0.04 *k*_*B*_*T*. Our final estimates of ΔΔ*G* are averages across predictions from converged replicates, with uncertainties propagated as 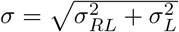. Our full protocol is described in Methods.

### 2.2 Convergence and uncertainty of expanded ensemble ΔΔ*G* estimates

The full set of EE calculations (113 mini-protein residues, each mutated to the other 19 canonical amino acids, in replicates of 10 for a total of 42940 L and RL simulations) provides an excellent opportunity to survey the factors that affect the predicted uncertainty of the ΔΔ*G* estimates. The distribution of estimated uncertainties is bimodal, with a major population centered near *σ* = 1.2 kcal/mol, and a minor population near *σ* = 6 kcal/mol (Figure 3a). The mean estimated uncertainty is 2.06 kcal/mol, with 66% of the estimates falling below that value.

**Fig. 3:**
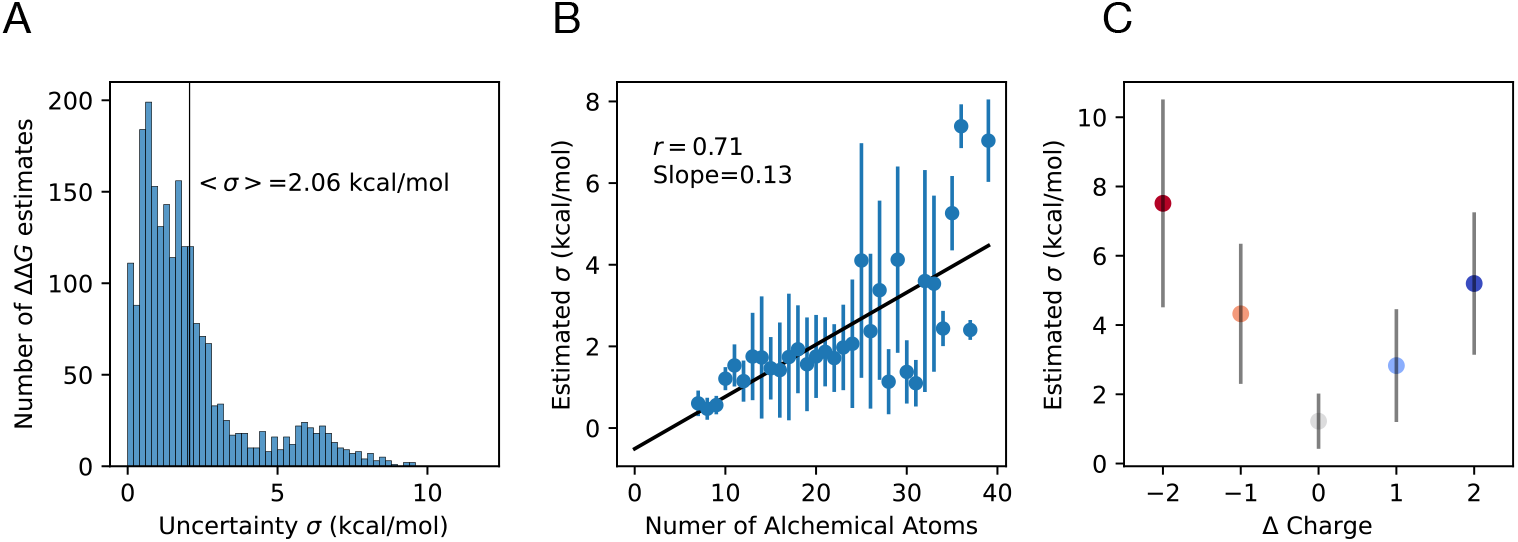
Estimated uncertainties of EE ΔΔ*G* estimates. (A) Histogram of estimated uncertainties, *σ*, for all 2147 ΔΔ*G* estimates. (B) Mean uncertainty estimates grouped by the number of atoms involved in the alchemical transformation. Error bars show standard deviations in each group. (C) Mean uncertainty estimates grouped by net charge change involved in the alchemical transformation. Error bars show standard deviations.

To further dissect the key factors affecting estimated uncertainties, we grouped the uncertainty estimates by various properties. Grouping the uncertainty estimates by the number of atoms involved in the alchemical transformation clearly shows strong correlation between the estimated uncertainty and the number of transformed atoms (Figures 3b, S1). These results suggest that up to 25 atoms can be alchemically transformed while still achieving average uncertainty estimates below 2 kcal/mol; beyond that number, EE estimates become significantly more unreliable.

Grouping the uncertainty estimates by the net change in amino acid charge reveals that largest uncertainties can be ascribed to transfomrations involving residues of different charges (Figures 3c, S2). While net-neutral transformations have average estimated uncertainties of about 1.2 kcal/mol, free energy estimates for mutations with ±1 and ±2 charge have average uncertainties of 3.8 kcal/mol and 6.4 kcal/mol, respectively. This suggests that relative free energy estimates for charge changes remains challenging, even with neutralizing counterions in the alchemical topology.[11, 28]

#### 2.2.1 Slow side chain dynamics correlates with increased uncertainty of EE free energy estimates

Another source of uncertainty comes from sampling bottlenecks when there are slow conformational degrees of freedom, especially those that occur on timescales close to the adaptive learning rate of the flat-histogram method used to obtained converges free energy estimates. While these bottlenecks can arise from complex alchemical transformations involving many atoms or charge changes, they can also arise simply from the slow molecular motions present in protein dynamics generally – for example, slow transitions between side chain rotamers in sterically hindered or otherwise rugged landscapes. Often these slow motions are coupled to solvent molecules that must enter or exit regions as the alchemical transformation proceeds.[29, 30]

To detect and quantify slow side chain dynamics in a systematic way, we analyzed the time-evolution of the side chain *χ*_1_ dihedral angle in each simulation (defined in all alchemical topologies except for G2A and A2G). The *χ*_1_(*t*) trajectory was discretized to a three-state space according to bins centered around (−60^*°*^, +60^*°*^, 180^*°*^), which was then used to build a simple three-state kinetic model using standard Markov State Model (MSM) approaches. A procedure was used to robustly estimate the slowest implied timescale from the ten simulation replicates (see Methods). This estimate corresponds to the correlation time of *χ*_1_-angle dynamics.

By grouping the results by amino acid type, we observe a roughly linear relation (Pearson *r*=0.34) between the implied timescale (ITS) of side chain dynamics and the uncertainty *σ* of the free energy estimate (Figure 4a). Charged/polar amino acids (E,K,N,Q,D) have slightly uncertainties, but trend similarly with the ITS. An outlier with small ITS but large uncertainty is arginine (R), a residue a net charge and a large side chain. Outliers with large ITS (larger than 150 ns) and small uncertainties include cysteine (C), which is involved in many interior disulfides (Figure 4a,b) and histidine (H) (Figure 4a,c).

**Fig. 4:**
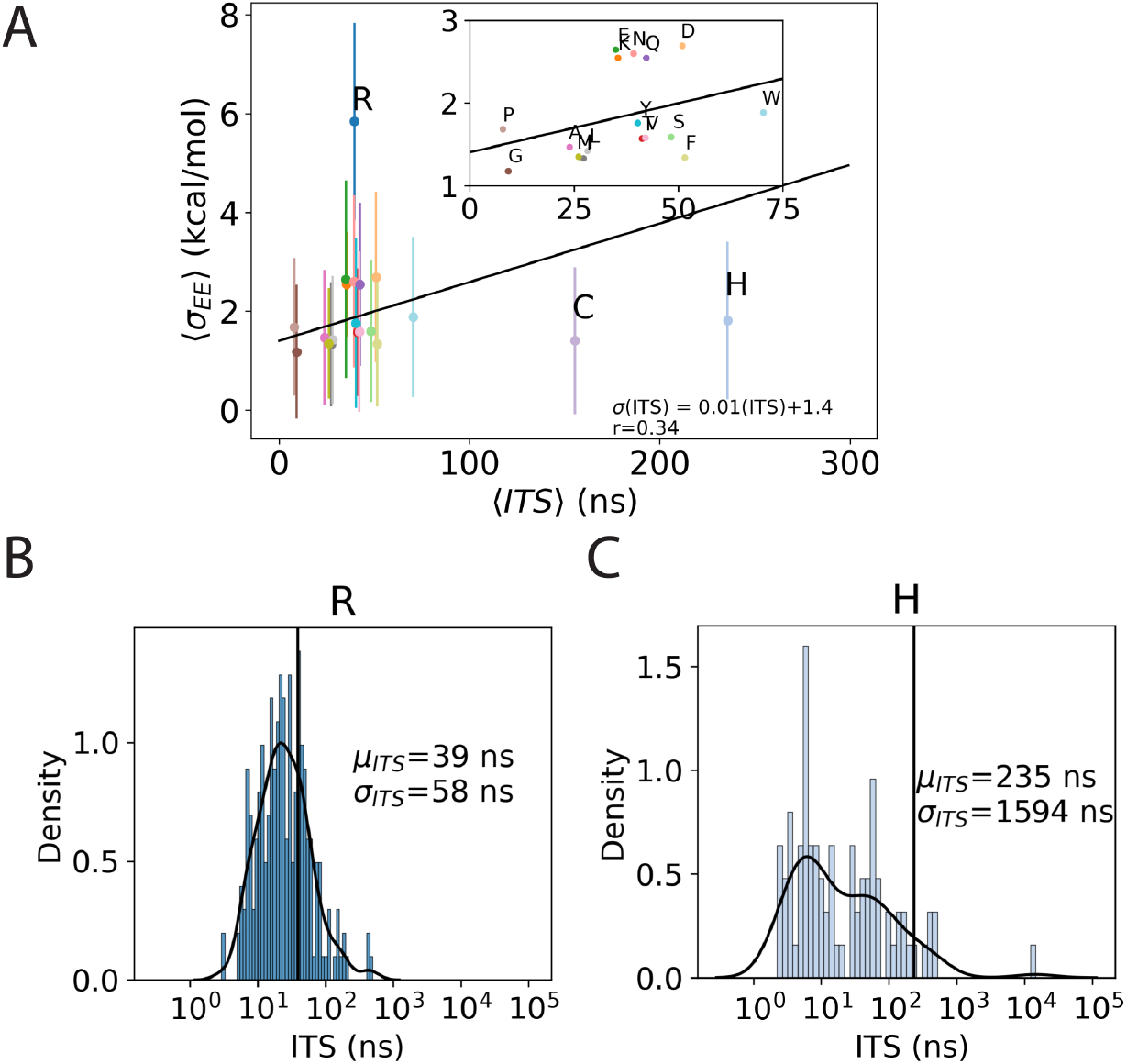
Implied timescales (ITS) of sidechain *χ*_1_-angle dynamics, grouped by amino acid type. (A) Mean implied timescales for each residue type plotted against mean uncertainty of FEP estimation. The linear regression line for non-outliers (line has slope of 0.1 kcal mol^−1^ ns^−1^ and intercept 1.4 ns) is shown in black, along with Pearson’s correlation coefficient (*r*=0.34). The inset shows a closeup of the non-outliers, labeled by amino acid. Error bars reflect the standard deviation. Histograms of implied timescales for (B) simulations with arginine (R), and (C) simulations with histidine (H). Kernel density estimates of the distributions are shown as a black line.

### 2.3 Bayesian inference of experimental ΔΔ*G* from high-throughput sequencing data

To compare FEP predictions of ΔΔ*G* to experiment, we developed a Bayesian method to infer *K*_*d*_ values from the high-throughput sequencing data published in Chevalier et al.[4] In those experiments, fluorescence-activated cell sorting (FACS) was used to select HA-bound yeast cells displaying designed miniprotein binders for next-gen sequencing. Sequencing with performed at a series of decreasing HA concentrations (1000 nM, 100 nM, 10 nM and 1 nM), and the affinity for a given miniprotein sequence was roughly characterized by the concentration at which the fraction of total sequence reads (the fraction enrichment) was maximal.

To infer *K*_*d,i*_ values for each miniprotein *i*, we make several simplifying assumptions. We assume the initial yeast pool are displaying the binders with efficiencies proportional to the numbers of sequence reads without HA selection. We also assume that fraction bound for each miniprotein depends on HA concentration like a single-species binding curve. By Bayes’ theorem, the posterior probability of the set of dissociation constants given the data *D* is

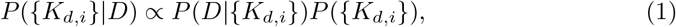

where *P* (*D*|{*K*_*d,i*_}) is the likelihood of observing the data *D* (i.e. the number of sequence reads for each sequence at all HA concentrations) given a set of *K*_*d,i*_ values, and *P* ({*K*_*d,i*_}) is a prior distribution. To model the likelihood function, we use a multinomial distribution conditioned on predicted bound fractions. As a model of the prior distribution *P* ({*K*_*d,i*_}), we fit a set of 1102 ΔΔ*G* values compiled from 57 protein-protein complexes in the SKEMPI 1.1 database by Geng et al.[31] to a Cauchy distribution, and model the prior as 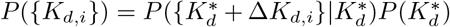, where 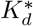 is the unknown dissociation constant wild-type miniprotein binder (A8, A13 or A18), which we treat as a nuisance parameter, modeling 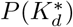 as a non-informative Jefferys’ prior.

To sample the posterior distribution, Markov Chain Monte Carlo (MCMC) was used in two rounds of 1M steps to converge estimates from initial starting values of 1 *μ*M (see Methods). After convergence, a final round of 10M steps was used for estimating mean values and uncertainties using the sample variance (Figure 5a). Large inferred *K*_*d*_ values have large uncertainties, because there are fewer sequence reads to restrain the estimates (Figure 5b). In all MCMC sampling, *K*_*d,i*_ values were constrained to be between 1 nM and 1 mM, as relative enrichment fractions provide little information to restrain absolute affinities. Predicted relative affinities are more robust; these are the important quantities for comparison with experimental SSM screens.

**Fig. 5:**
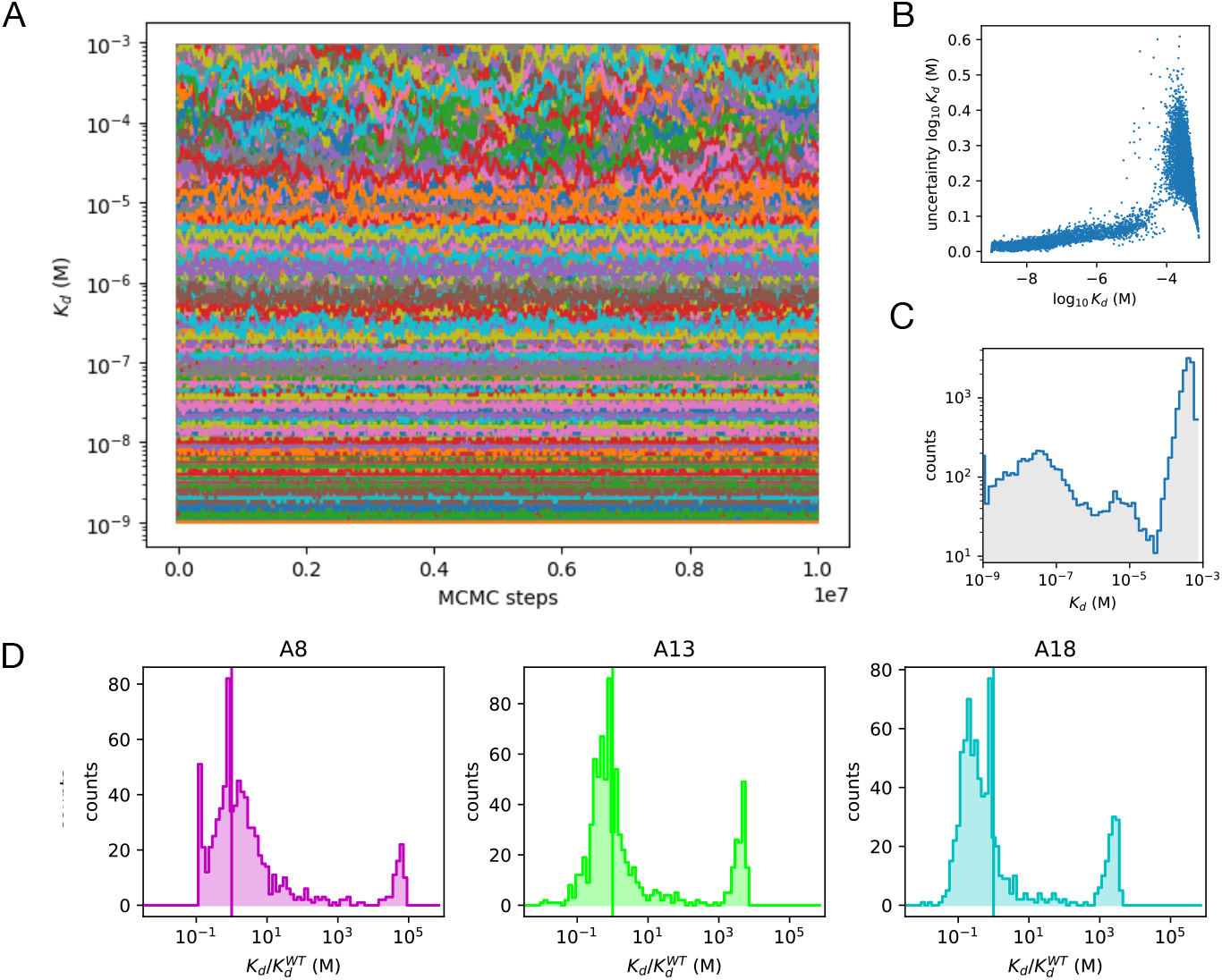
Bayesian inference of miniprotein binding affinities from high-throughput sequencing data. (a) Traces of *K*_*d*_ estimates for all 15490 sequences in the high-throughput sequencing data, throughout 10M steps production-run MCMC sampling of the Bayesian posterior. (b) Scatter plot of the estimated uncertainty in the log_10_ *K*_*d*_ of each sequence, versus log_10_ *K*_*d*_. The tapering at large log_10_ *K*_*d*_ values is due to the sampling upper bound of *K*_*d*_ = 1 mM. (c) The distribution of the 15490 inferred *K*_*d*_ values. (d) Distributions of inferred relative 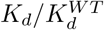 values for A8, A13, and A18 SSM sequences.

The published sequencing data for the affinity-maturation experiments comprise 15490 sequences, only 2394 of which correspond to A8 (796), A13 (801) and A13 (797) SSM variants. For the other 13096 miniprotein sequences in the published dataset, most have inferred *K*_*d*_ values in the millimolar range (Figures 5c). For the SSM sequences, most of the inferred *K*_*d*_ values range within an order of magnitude from the wild-type sequence, while a smaller population have inferred *K*_*d*_ more than two orders of magnitude larger than the wild-type, indicating a deleterious mutation (Figure 5d).

### 2.4 ΔΔ*G* predictions from Rosetta

Since Chevalier et al. 2017, improvements in Rosetta-based searching and scoring have been made,[19, 32–35] although optimizing binding using purely *in silico* mutational scanning remains challenging. [36–39] To compare our *in silico* SSM ΔΔ*G* predictions to current state-of-the-art Rosetta predictions, we used RosettaDDGPrediction, a tool that facilitates high-throughput mutational scanning using a number of Rosetta protocols and scoring functions.[40] We used the Flex ddG protocol[19]) with the *talaris2014* [41] scoring function, a combination previously shown to achieve mean unsigned errors near 1 kcal/mol across a large (N = 1240) test set.[19]

### 2.5 Comparison of ΔΔ*G* predictions from *in silico* SSM screening with experiment

The complete set of SSM ΔΔ*G* predictions from EE and Flex ddG, and the inferred experimental ΔΔ*G* values, are shown as a set of heatmaps, using a red-to-blue color scale where stabilizing mutations (ΔΔ*G <* 0) are shown in shades of red and destabilizing mutants (ΔΔ*G >* 0) are shown in shades of blue (Figure 6). Visually, the EE results have a pattern similar to that of the experimental values, with residues positions in the conserved binding helix motif showing positive ΔΔ*G* values for all (or most) mutations. The Flex ddG predictions also show this pattern, but with a smaller variance.

**Fig. 6:**
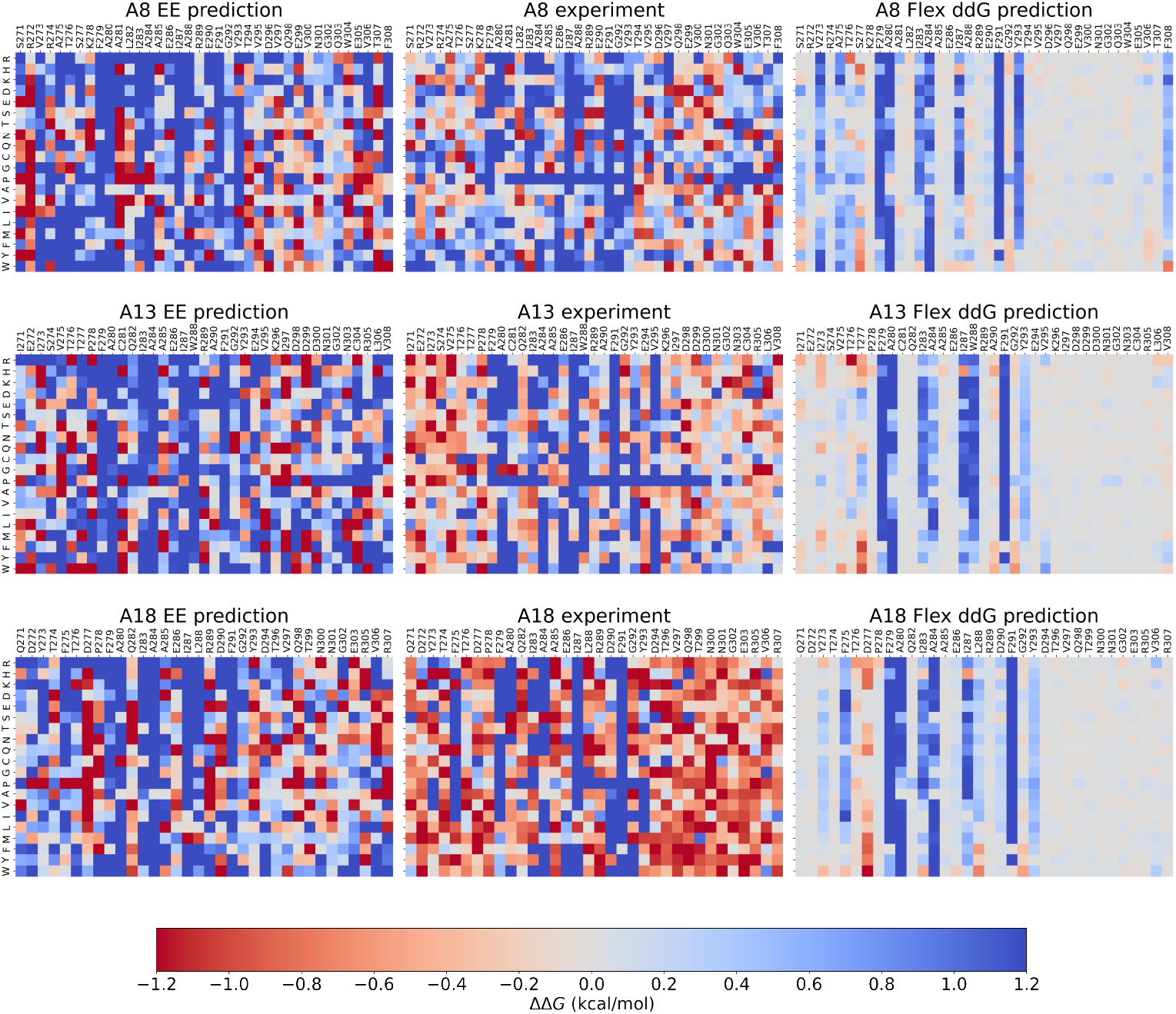
Site Saturated Mutagenesis (SSM) Heatmaps. Left column: Predicted changes in binding free energy ΔΔ*G* from expanded ensemble free energy calculations. Center column: Experimental ΔΔ*G* values inferred from high-throughput sequencing data from Chevalier et al (2017). Right column: Predicted ΔΔ*G* values from the Flex ddG Rosetta protocol with the *talaris2014* scoring function.

Scatter plots of predicted vs. experimental ΔΔ*G* for the complete set of 2337 SSM variants show that EE free energy estimates are generally predictive for all three designed miniproteins (Pearson *r* ≈0.42), although there is a subset of predictions for destabilizing mutations with large outliers (ΔΔ*G >* 7.5 kcal/mol) (Figure 7). As a result, the mean unsigned error (MUE) of the EE ΔΔ*G* predictions exceed 2.0 kcal/mol.

**Fig. 7:**
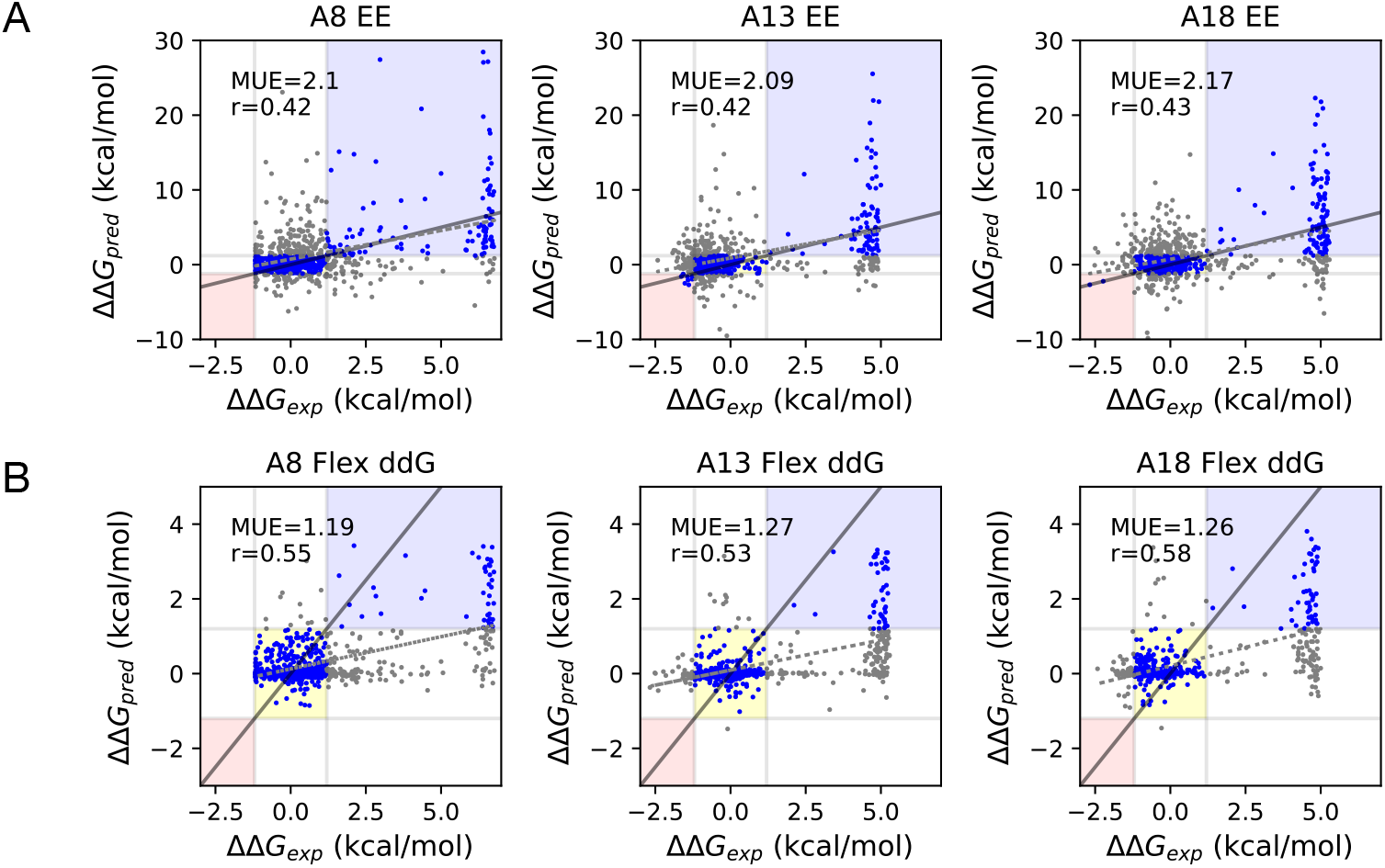
Comparison of predicted and inferred experimental values of ΔΔ*G*. Scatter plots of (A) expanded ensemble (EE) predictions of ΔΔ*G*, and (B) Flex ddG predictions of ΔΔ*G*, versus experimental values for A8, A13 and A18. Plots are annotated with the mean unsigned error (MUE, kcal/mol) and the Pearson correlation coefficient, *r*. The black line denotes unity (exact agreement), while the dashed gray line denotes the best linear fit. Shaded regions denote correct classification of stabilizing (pink), neutral (yellow), and destabilizing (blue) mutations, defined using a threshold of *±*1.2 kcal/mol.

In comparison to the EE estimates, Flex ddG predictions of ΔΔ*G* are generally more accurate (MUE *<* 1.27 kcal/mol, *r >* 0.53), but more conservative. Whereas linear regression fits of the EE predictions to experiment have slopes above 0.74, fits of the Flex ddG predictions give slopes ∼ 0.2 (Figure 7, Table 1).

**Table 1:** Accuracy of ΔΔ*G* predictions across the set of 2337 SSM variants.

| Method | miniprotein | MUE (kcal/mol) | Slope | r |
| --- | --- | --- | --- | --- |
| EE | A8 | 2.1 | 0.77 | 0.42 |
|  | A13 | 2.1 | 0.74 | 0.42 |
|  | A18 | 2.2 | 0.74 | 0.43 |
| Flex ddG | A8 | 1.2 | 0.17 | 0.55 |
|  | A13 | 1.3 | 0.16 | 0.53 |
|  | A18 | 1.3 | 0.20 | 0.58 |

### 2.6 Classification of stabilizing and destabilizing mutations

We next asked how well ΔΔ*G* estimates from EE and Flex ddG were able to correctly predict stabilizing and destabilizing mutations, which we define as ΔΔ*G ≤* −1.2 kcal/mol and ΔΔ*G*≥ +1.2 kcal/mol, respectively, following previous conventions in the literature.[40, 42] We consider a mutation to be ‘neutral’ otherwise. With these classification categories, the confusion matrix of classification errors shows that Flex ddG is highly conservative, predicting all stabilizing mutations to be neutral (Figure 8a). EE predicts 235 mutations to be stabilizing, correctly identifying 12.

**Fig. 8:**
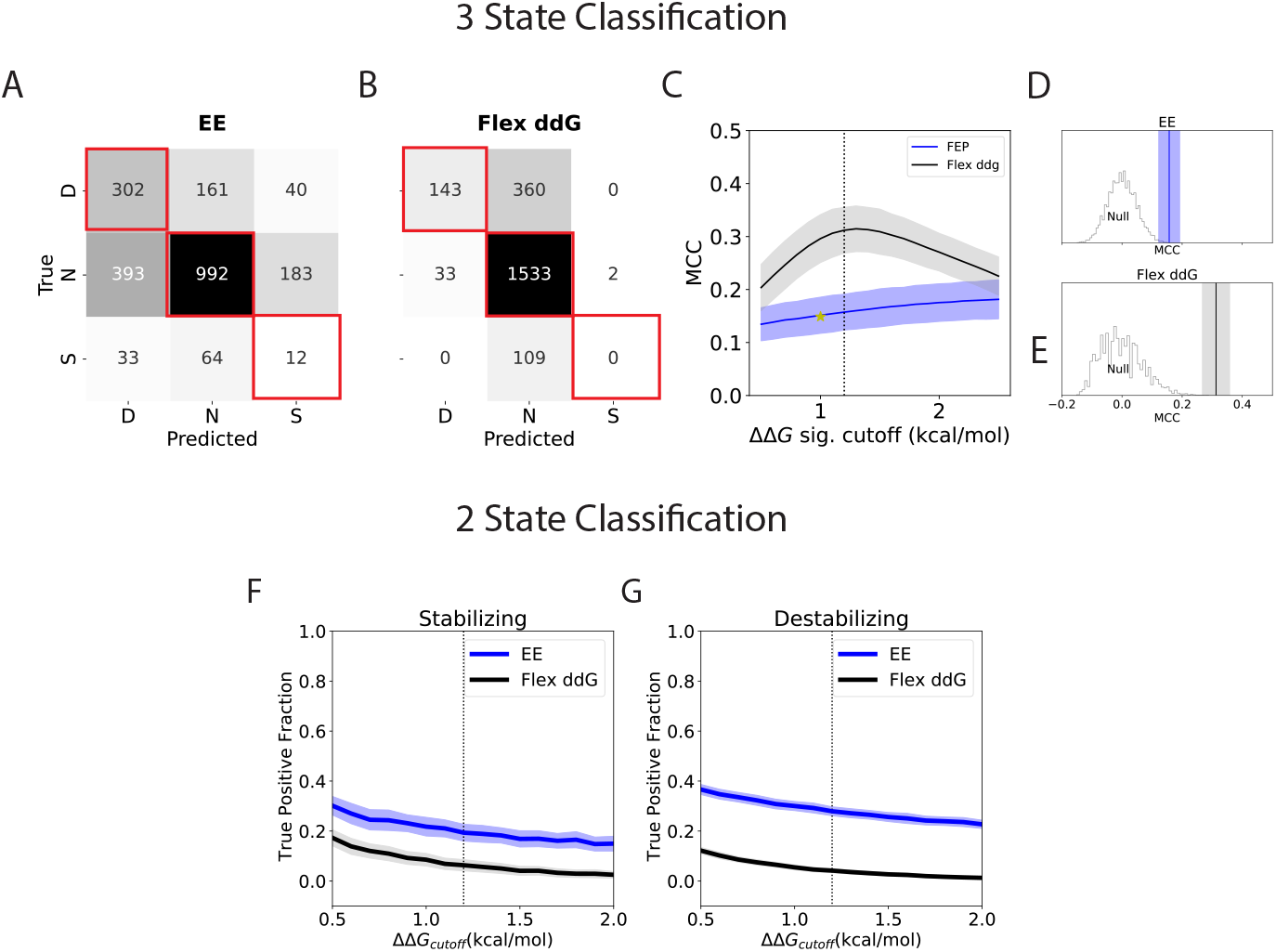
Classification of stabilizing and destabilizing mutations: performance for EE and Flex ddG. Confusion matrices for (A) EE and (B) Flex ddG, displaying the numbers of predicted vs. actual classifications of destabilizing (D), neutral (N), or stabilizing (S) mutations. (C) Sensitivity of the MCC to the significance cutoff used in the prediction classifier. Vertical line is shown at the ±1.2 kcal/mol threshold. Shaded regions represent MCC uncertainties propagated from predicted ΔΔ*G* uncertainties, calculated using a bootstrap procedure. Gold star marks the MCC of 0.15 for the combined dataset analyzed in Cao et al. 2022. Null distributions of the MCC constructed from random permutations of classification labels for (d) EE and (e) Flex ddG show that the computed values of MCC using the ±1.2 kcal/mol threshold are statistically significant. True positives rates for two-state classification of stabilizing and destabilizing mutations are shown for (f) EE and (g) Flex ddG, as function of the significance cutoff used in the classifier.

The majority of mutations in the experimental SSM scans are neutral (1568); 503 mutations are destabilizing, and only 109 are stabilizing. Because of this, true positive rates alone are misleading; for example, simply predicting that all mutations are neutral would achieve a true positive rate of 72%. To better quantity the success of each method’s ternary classification of stabilizing, neutral, and destabilizing mutations, we use the Matthews Correlation Coefficient (MCC), which has been shown to perform robustly on unbalanced datasets for both binary and multi-class classifiers.[43] The MCC is defined as

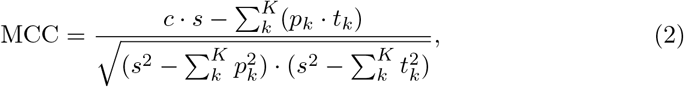

where 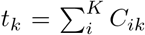 is the number of times class *k* truly occurred, 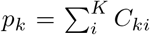 the number of times class *k* was predicted, 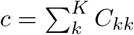 the total number of samples correctly predicted, 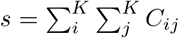 the total number of samples.

Using a |ΔΔ*G*| ≥ 1.2 kcal/mol significance cutoff, the MCC for the Flex ddG predictions is 0.31± 0.05, while the MCC for the EE predictions is 0.16 ± 0.03. To probe the sensitivity of the predictions to the significance cutoff, we varied the cutoff for the predicted classifications from 0.5 to 2.5 kcal/mol, while keeping the threshold for experimental significance at 1.2 kcal/mol (Figure 8c). As the significance cutoff increases, the MCC for Flex ddG decreases and the MCC for EE increases, with a cross-over point beyond 2.5 kcal/mol. Construction of the null distribution of the MCC statistic for EE and Flex ddG by random permutation of classification labels confirms that predicted classifications are statistically significant (*p <* 4 ×10^−4^, Figure 8d,e).

We also examined results from a more recent study by Cao et al. (2022) which measured experimental ΔΔ*G* for 12 *de novo* designed miniprotein binder SSMs and compared against predictions from Rosetta.[3] This dataset comprises 117728 comparisons, grouped into five structural categories: interface core (13298), interface boundary (25162), monomer core (18875), monomer boundary (19138), monomer surface (41255). Using the authors’ classification threshold of 1.0 kcal/mol, the computed MCC for these predictions range from 0.09 to 0.21, with an MCC of 0.15 for the combined dataset.

How accurate are EE and Flex ddG at simply predicting stabilizing or destabilizing mutations? To answer this, we computed true positive rates for stabilizing mutations (Figure 8f) and destabilizing mutations (Figure 8g), across a range of significance thresholds. In this case, we find that EE outperforms Flex ddG, suggesting that the large MCC values for Flex ddG are primarily due to its conservative classification of neutral mutations. A similar analysis for the Cao et al. 2022 results show true positive rates of stabilizing mutations similar to Flex ddG (0.13), and surprisingly good true positive rates for destabilizing mutations (0.77, see Figure S6).

### 2.7 Prediction of average residue fitness

To assess the extent to which EE and Flex ddG can predict average mutational effects at each position, we computed the average ΔΔ*G* across all amino acid substitutions, for each position (Figure 9a). Linear regression fits to the inferred experimental averages show mean unsigned errors within 1 kcal/mol for both EE (0.95 kcal/mol) and Flex ddG (0.79 kcal/mol), each with similar correlation (Pearson’s *r* of 0.72 for EE and 0.77 for Flex ddG). While the slope of the EE regression is nearly unity, the slope of the Flex ddG regression is 0.31, indicating the conservative nature of Flex ddG predictions for average ΔΔ*G*, which have a narrower distribution than EE or the inferred experimental values (Figure 9b). Predictions of average ΔΔ*G* mapped to the three-dimensional structure of each miniprotein are shown in Figure 9c.

**Fig. 9:**
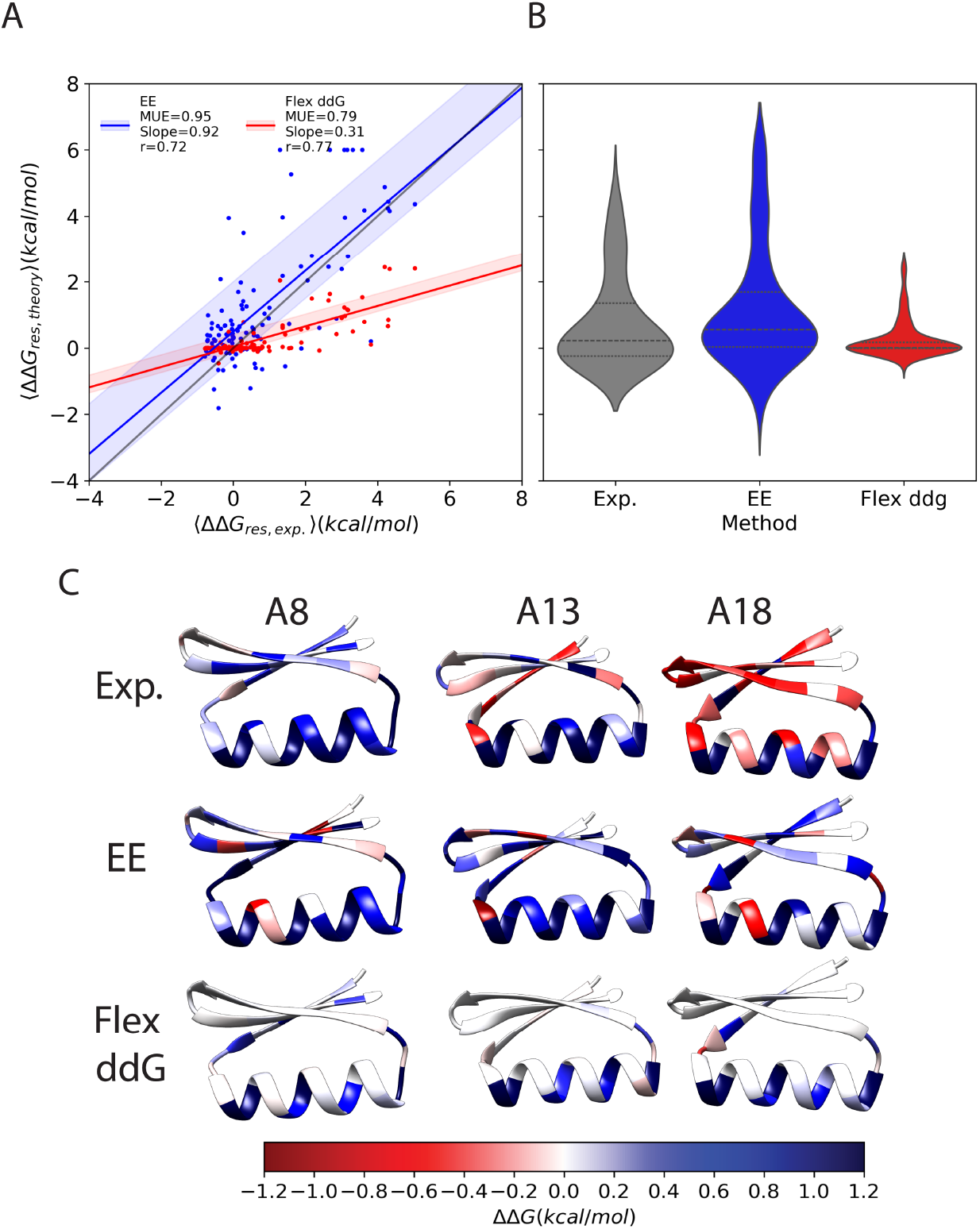
Prediction of average mutational effects at each position. (A) Scatter plot of average predicted ΔΔ*G* for EE (blue) and Flex ddG (red) at each position versus the average inferred experimental values. Solid lines are linear regression fits, with shaded regions corresponding to the the 95% confidence interval. The legend shows the mean unsigned error (MUE), slope, and Pearson *r*-value for each regression. The line of unity is shown in gray. (B) Violin plots showing the distribution of average mutational effects for experiment (gray), EE predictions (blue) and Flex ddG predictions (red). (C) Structures of miniproteins A8, A13 and A18 colored at each position by average ΔΔ*G*: (top) inferred experimental values, (center) predictions from EE, and (bottom) predictions from Flex ddG.

### 2.8 Can SSM predictions rationalize observed affinity matured variants?

While miniproteins A8, A13 and A18 each bind HA with a 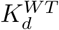 above 300 nM, the affinity-matured variants each contain five, six, or seven mutations which together confer low nanomolar affinity, with 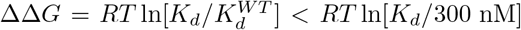. This implies that the Δ*DeltaG* values for the affinity-matured variants have an upper bound of -2.3 kcal/mol (Table 2). To what extent can single-residue ΔΔ*G* values rationalize these sets of mutations?

**Table 2:** Affinity Maturation Mutations of HA Binders.

| Miniprotein | Mutations | $\Delta\Delta G$ (kcal/mol) |
| --- | --- | --- |
| A8 | R272G, V300K, Q303E, E305R, T307V | $< -2.32$ |
| A13 | I273Q, V275F, P278L, I283T, E294T, D298M, D299V | $< -2.73$ |
| A18 | D272E, T274R, D277N, A285L, D294A, N300I | $< -2.97$ |

A simple additive model of the effects of multiple mutations, ΔΔ*G*_affinty-matured_ = Σ_*i*_ ΔΔ*G*_*i*_, where ΔΔ*G*_*i*_ are the inferred experimental values, can satisfactorily explain maturation of A13 and A18, but not A8 (Figure 10). Interestingly, T307V is the only single-residue mutant with increased affinity, which suggests strong epistatic effects.

**Fig. 10:**
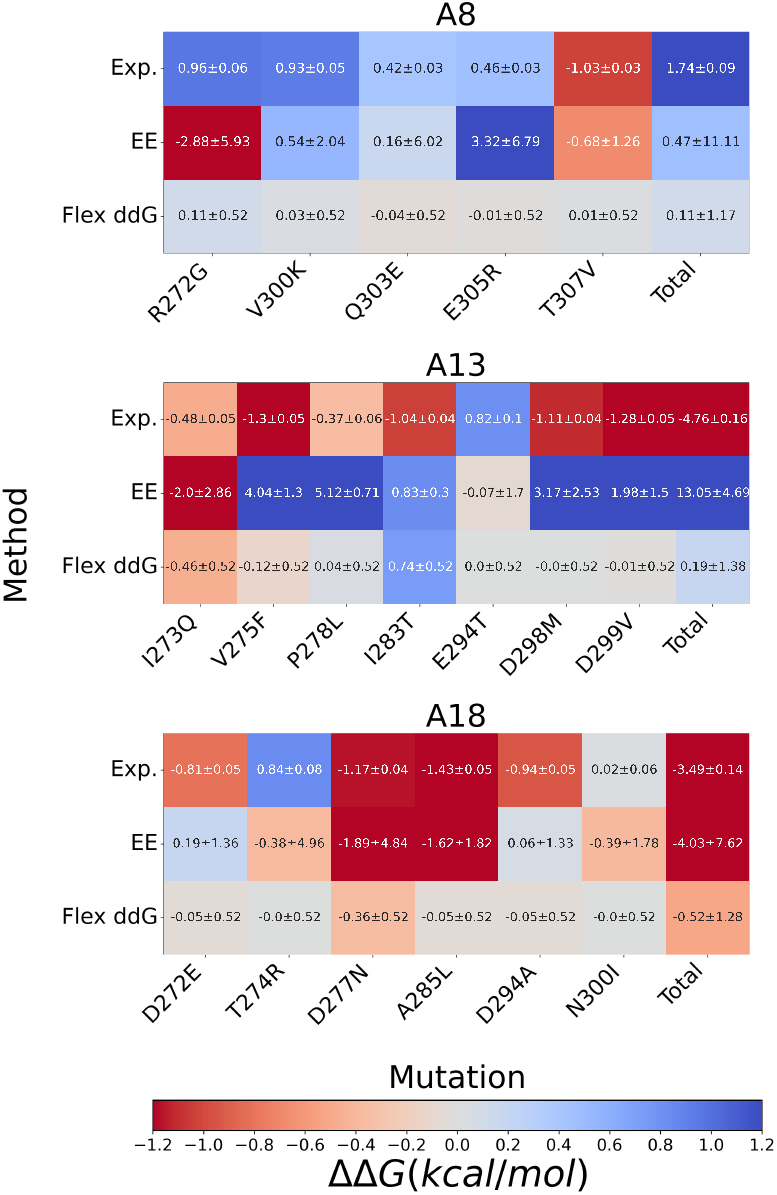
Additive effects of single-residue mutations cannot fully explain observed affinity-matured variants. (Top) Heatmaps of ΔΔ*Gs* for specific single-residue mutations observed in affinity-matured miniproteins. Each column is a mutation, with the rightmost column showing the additive total of all single-residue mutations. Shown for each miniprotein are inferred experimental values (Exp.), expanded ensemble predictions (EE), and predictions from Flex ddG, along with estimated uncertainties.

When predictions of ΔΔ*G*_*i*_ are used with the additive model, EE predicts only one of the miniprotein variants to have higher affinity, and with large uncertainties. Flex ddG predicts two of the three variants to have higher affinity, but only weakly (ΔΔ*G* ≈ −0.5 kcal/mol), as single-residue ΔΔ*G*_*i*_ predictions are consistently conservative (|ΔΔ*G*_*i*_| *<* 0.46 kcal/mol in all cases) Overall, single-residue mutations do not provide a clear picture of the mechanism of affinity maturation for the three variants.

### 2.9 Can simple electrostatic models rationalize sequences with increased affinity?

The non-additivity of single-residue mutations suggests epistatic effects play a role in the affinity maturation of miniproteins A8, A13 and A18. A simple physical hypothesis is that electrostatic interactions are important, especially considering four of the five affinity maturation mutations to A8 involve ionizable residues.

To test the importance of electrostatics, we consider two simple metrics of electrostatic interaction that depend on the sequence *s*, conditioned on the structural model *M*. The first metric,

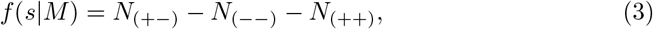

counts the difference between favorable and unfavorable salt-bridges. This is calculated by constructing a contact map of heavy atom side chain contacts of *<*= 4.5*Å* both intra-miniprotein and with HA (see Tables S1–S3 and Supplementary Fig. S7-S9)

The second metric, *c*(*s* |*M*), is based on a graph-theoretic quantity called the Laplacian centrality.[44] We construct a protein contact graph *G* with edges weighted *w* = 1 if the contact is electrostatically repulsive, *w* = 2 if neutral, or *w* = 3 if attractive. The Laplacian centrality *CL*_*i*_ = Δ*E*_*i*_*/E*_*L*_(*G*) is defined for a node *i* (residue position) in the graph; it is the fractional difference in the Laplacian energy after removing node *i* from the graph, where the Laplacian energy 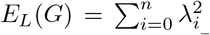 is the sum of the eigenvalues of the graph Laplacian matrix. The metric *c*(*s*|*M*) = Σ_*i*_ *CL*_*i*_ is the sum across all residue positions.

To test whether the electrostatic interactions quantified by these metrics are important, we used a simulated tempering algorithm to sample the Boltzmann distribution of all possible 5- or 6-residue mutants assuming an additive model of affinity maturation, and compared these results to a null model where each sequence is equally likely (i.e. infinite temperature) (Figure S10). In each case, we found that the distributions of *f* (*s*|*M*) and *c*(*s*|*M*) (Figure S11) are not significantly different from the null distribution, suggesting that simple models of electrostatics (i.e. quantifying salt bridge interactions) provide an incomplete picture of the fitness landscape.

### 2.10 Affinity matured variants have mutations at highly mutable positions

Using the simulated tempering (ST) sampling described above, we compute for each miniprotein the Shannon entropies at each position from the observed frequencies of each amino acid (Figure 11). As expected, the Shannon entropy profiles show low sequence entropy for residues in the conserved helix binding motif (e.g. F279, I287, F291). Residue positions that are mutated in the affinity matured variants have high sequence entropy. To visualize these values on the three-dimensional structures of each miniprotein, we compute a “mutability” score by normalizing the Shannon entropies to unit variance and subtracting the mean. Nearly all of the mutations in the affinity matured variants are found at highly mutable positions.

**Fig. 11:**
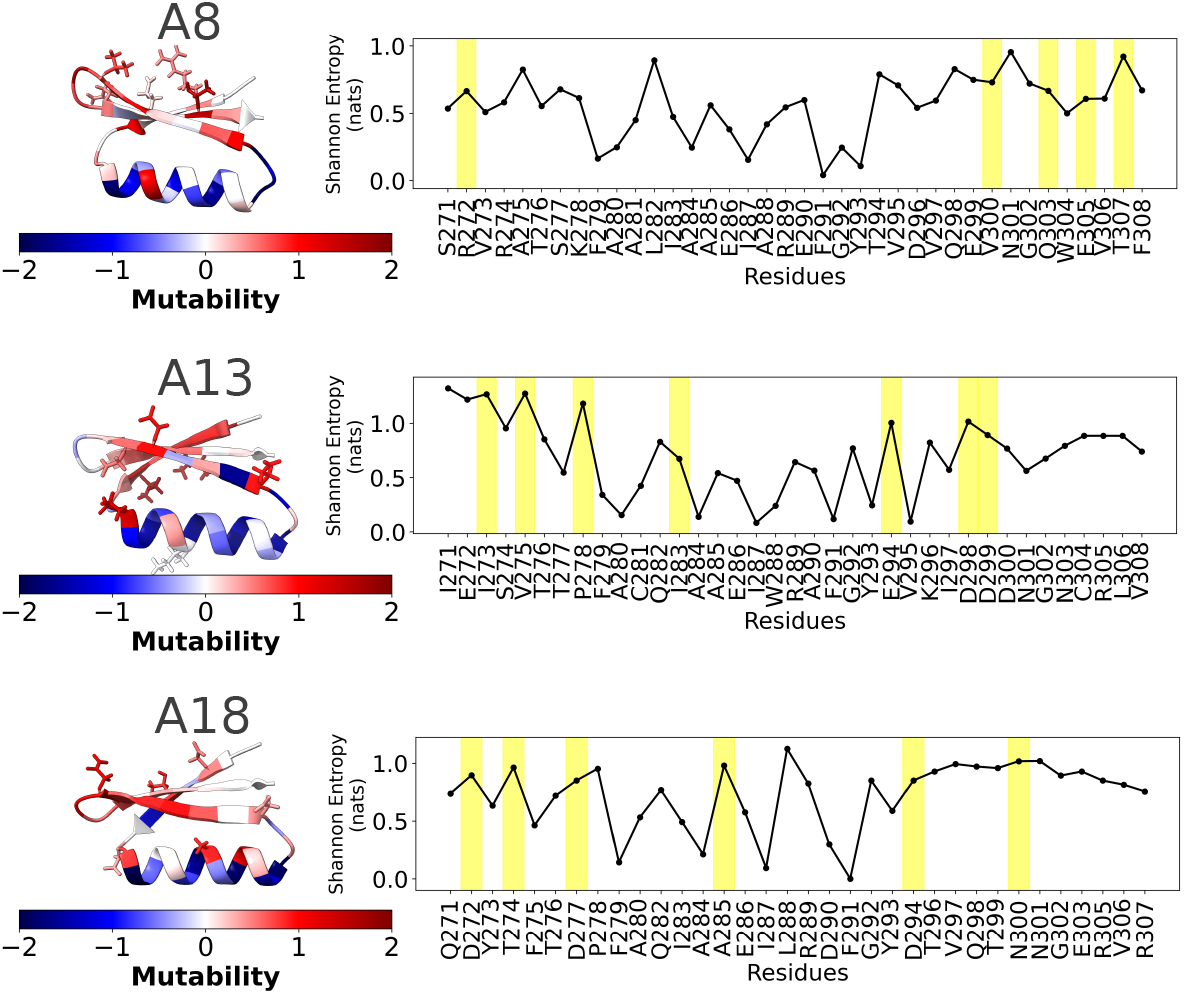
Mutability profiles of designed miniproteins. Shannon entropies computed from inferred experimental frequencies *e*^−ΔΔ*G/RT*^ are shown for each miniprotein residue position. Mutability scores (mean-subtracted Shannon entropies normalized to unit variance) are shown color-mapped to the three-dimensional structure of each miniprotein. Mutations observed in the affinity-matured variants (yellow bars) often coincide with positions having high mutability.

Shannon entropy profiles computed from the ST sampling are highly similar to those computed from position-independent Boltzmann weighted frequencies 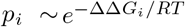. Shannon entropies computed using EE predictions of ΔΔ*G*_*i*_ show similar profiles, while those computed using the more conservative Flex ddG predictions show higher entropies with less variation (Figure S12).

## 3 Outlook

To our knowledge, this is the largest dataset of expanded ensemble simulations constructed to date, as well as the largest set of relative binding free energy calculations performed for independent constructs. It adds to the recent interest in and extension of the expanded ensemble method, which has distinct advantages over other existing approaches due to its asynchronous parallelizability. An assessment of the quality of free energy methods such as EE for large-scale mutational prediction is timely, as many new methods are being developed to examine protein fitness landscapes through the combination of free energy approaches and machine learning.[11, 13, 45]

While our simulation-based EE protocol is more computationally expensive than algorithms like Rosetta, our work suggests it is better at identifying stabilizing mutations, a primary goal of protein design. Further improvements in accuracy are likely within reach that can justify this expense. For example, our work and the work of others have identified sampling barriers–both in conformational motions and across thermodynamic ensembles–as a major source of uncertainty in free energy calculations.[16, 29] Recent successes in identifying such barriers and designing enhanced sampling schemes to address them,[24, 46] suggest ways to improve sampling. For EE in particular, better adaptive sampling schemes beyond flat-histogram methods are also likely to improve the uncertainties of free energy estimates.[47–50] Moreover, the EE calculations presented here were performed using CPU-enabled GROMACS; further efficiencies from GPU acceleration is an obvious and highly feasible next step. The accuracy of massively parallelizable physical free energy approaches like EE will also likely benefit from continuing advances in deep learning. Already, a hybrid neural-network potential /molecular mechanics energy function shows best-in-class accuracy for a benchmark set of relative binding free energy calculations.[51] Deep learning approaches have also made amazing strides towards automated protein design in the last few years.[52–54] Augmenting deep methods with physics-based approaches like EE is a particularly exciting prospect, as physical models can access much large chemical spaces beyond the canonical twenty amino acids, as well as modeling functional dynamics.

## 4 Conclusion

In this work, we use an expanded ensemble (EE) approach to perform a massively parallel set of relative binding free energy (RBFE) calculations for site-saturated mutants (SSM) of three *de novo* designed minprotein binders to hemaggluttinin. As far as we are aware, these calculations are the most comprehensive alchemical FEP datasets for designed protein-protein interactions, encompassing nearly 50K individual simulations. We compared predicted changes in binding free energies from EE to experimental values inferred from published high-throughput sequencing data using a Bayesian inference method. We additionally assessed predictions from the Flex ddG Rosetta protocol.

The results show that EE methods can—from structural models alone—predict changes in binding affinity within about 2.1 kcal/mol on average. Systematic comparisons identify several contributions to large uncertainty: net charge changes, large numbers of atoms involved in alchemical transformations, and slow conformational dynamics in residue side chains. Flex ddG predictions are on average more accurate (about 1.2 kcal/mol mean unsigned error), but this largely reflects the conservative prediction that most mutations are neutral. In contrast, EE can better classify stabilizing and destabilizing mutations. We also explored the ability of SSM scans to rationalize known affinity-matured variants containing multiple mutations, which cannot be explained from additive models, likely due to epistatic effects. Although simple electrostatic models fail to explain nonadditivity, we show that observed mutations are found at more mutable positions having higher Shannon entropies.

Overall, this study suggests that simulation-based free energy methods can provide valuable and predictive information for *in silico* affinity maturation of designed miniproteins, with many feasible improvements to the efficiency and accuracy within reach.

## 5 Methods

### 5.1 Molecular Simulation

#### System preparation

Initial coordinates of miniproteins A8, A13 and A18 in complex with a truncated version of hemagglutinin containing HA1 and HA2 binding regions were taken from structural models published by Chevalier et al.[4]. Hybrid alchemical topologies for complex and monomer simulations were constructed using GROMACS 2020.3[20, 55] and *pmx* (https://github.com/deGrootLab/pmx),[22] using the *amber14sbmut* force field.[23] For transformations involving net charge changes, one or two neutralizing counterions were coupled to the alchemical topology and restrained at a position 1.0 nm from the solute. Preparation scripts were written in Python and utilized the mdtraj package.[56]

Systems were solvated with TIP3P water and 150 mM NaCl in a periodic box. Monomer systems comprised ∼14K atoms in a 5.3 nm cubic box; complex systems comprised ∼93400 atoms in a 9.82 nm cubic box. Position restraints were applied to the backbone N, C, and CA atoms of the miniprotein and HA, using an isotropic force constant of 1000 kJ mol^−1^ nm^−2^. During analysis we discovered that C281 and C304 were in their reduced form instead of a disulfide bond; we suspect only minor effects on our results since these residues are in the interior of the miniprotein, and are position restrained.

Systems were steepest-descent minimized and temperature equilibrated at 300 K for 100 ps using a leap-frog integrator. Production simulations were performed in the NVT ensemble using velocity Verlet integration with a 2 fs time step, hydrogens constrained using LINCS, PME electrostatics, 0.9 nm vdW cutoffs, and a long-range dispersion correction. All simulations were performed using GROMACS 2020.3 on the Folding@home distributed computing platform.[21]

#### Expanded ensemble protocol

The GROMACS implementation of EE[57] was used with 45 intermediates along the alchemical transformation. Soft-core potentials were used with *sc-alpha*=0.5, *sc-power* =1, and *sc-sigma*=0.3. Exchanges were proposed/accepted every 1 ps according to a metropolized-Gibbs criterion. The initial Wang-Landau (WL) increment was set to *δ* = 10 *k*_*B*_*T*. The histogram of sampled intermediates was deemed sufficiently flat when all histograms counts *h*_*i*_ come wihtin some threshold of the mean, 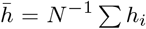, such that min 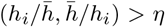 for all *i*, where *η* is set to 0.7. Upon reaching this criterion, the histogram counts are reset to zero and the WL increment is scaled by *α* = 0.8.

#### Optimization of λ-values

Intermediates spanning the alchemical transformation are parameterized by a series of *λ*_*i*_ values, *i* = 1, …*N*, ranging from *λ*_1_ = 0 to *λ*_*N*_ = 1. The choice of the *λ*_*i*_ is system-dependent, and very important to the convergence of EE estimates; poor choices produce low acceptance of proposed moves, resulting in slow convergence. To learn optimal *λ*_*i*_ schedules for each transformation, we first performed pre-production simulations using a “one size fits all” strategy: Initial values for non-charge changing transformations were:

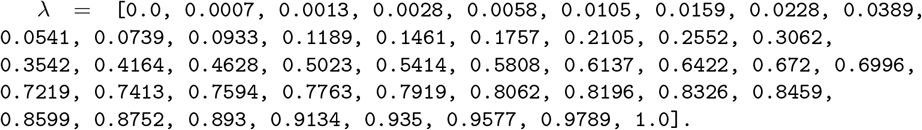

Initial values for charge changing transformations, based on a 20 ns test for A8 R272E, were set to:

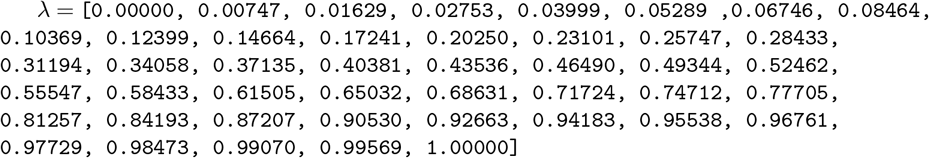

To refine these parameters for each system, EE sampling was performed on Folding@home for at least 20 ns, restricted to nearest-neighbor transitions in *λ*_*i*_ to ensure sampling of all alchemical intermediates. Written to file every 1 ps are the full set of reduced energy perturbations Δ*u*_*ij*_ = *β*(*U* (*x*_*i*_|*λ*_*j*_) − *U* (*x*_*i*_|*λ*_*i*_) where *x*_*i*_ is the current configuration being sampled in ensemble *i*, and *β* = 1*/k*_*B*_*T*. The *pylambdaopt* algorithm (https://github.com/vvoelz/pylambdaopt) is then used to infer improved values 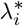 that minimize the approximate thermodynamic length[58, 59] between all pairs of neighboring intermediates. To second-order approximation, the thermodynamic length |*𝓁*(*λ*_*k*+1_)−*𝓁*(*λ*_*k*_)| between intermediates *k* and *k*+1 is proportional to the mean variance of the distributions *P* (Δ*u*_*k*(*k*+1)_) and *P* (Δ*u*_(*k*+1)*k*_) [60]. The *pylambdaopt* algorithm uses cubic spline fitting to first find a continuous differentiable function *𝓁*(*λ*) that interpolates the estimated *𝓁*(*λ*_*i*_). Then steepest-descent minimization is used to find new values 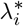 that minimize the loss function ℒ = Σ_*k*_ |*𝓁*(*λ*_*k*+1_) − *𝓁*(*λ*_*k*_)|^2^. The optimized 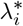 values were used for the remainder of each production run.

### 5.2 Implied timescales for sidechain dynamics

From trajectories of *χ*_1_ sidechain dihedral angles over time binned into trans, gauche+ and gauche-states, 3 × 3 transition count matrices *C*_*ij*_ (*τ*) were constructed of the number of sliding-count transitions observed from state *i* to state *j* within lag time *τ*. From the count matrix, Deeptime [61] was used to infer a Markov State Model (MSM) transition matrix *T*_*ij*_ (*τ*) using a maximum likelihood estimator with a constraint on detailed balance. The implied time scales (ITS) *t*_*n*_ at a lag time *τ* are computed as *t*_*n*_ = −*τ/*(ln *μ*_*n*_), for *n* = 2, 3, where *μ*_*n*_ are the eigenvalues of **T**.[62]

Typically, the MSM lag time is chosen from an inspection of an implied timescale plot, as the value of *τ* beyond which the time scales *t*_*n*_ plateau. For each alchemical transformation (for miniproteins bound to HA), a moving-block bootstrap procedure was used to sample count matrices across the ten EE trajectory replicates, at lag times ranging from 1/2 to 1 times time length of the shortest trajectory, to compute anaverage ITS curve and estimated uncertainties.

To calculate a standardized lag time to be used across all models, we numerically calculate the derivative of each ITS curve to find the lag time *τ* ^∗^ at which the slowest ITS (*t*_2_) plateaus. We calculate the mode of *τ* ^∗^ across all mutations at a given position (Figure S3), and then take the mean of the modes across all positions in the three miniproteins, which was ∼21 ns. Therefore, we used a standard lag time of *τ* =20 ns for all ITS analysis.

### 5.3 Flex ddG calculations

Relative binding free energies were calculated with RosettaDDGPrediction[40] using Rosetta 13.3 from the weekly release of Dec. 11 2023 (`rosetta.binary.ubuntu.release-362`). The Flex ddG protocol[19] was used with the *talaris2014* scoring function.[41] We selected the miniprotein as the protein to move, generating ten structures per mutation. An extra rotamer cutoff of 18 neighbors and 35000 backrub trials was used, with a stride of 7000. A modified version of `RosettaDDGPrediction/aggregation.py` was used to compile the prediction data.

### 5.4 Bayesian inference of dissociation constants from high-throughput sequencing data

To compare FEP predictions of ΔΔ*G* to experiment, we developed a Bayesian method to infer *K*_*d*_ values from the high-throughput sequencing data published by Chevalier et al.[4] Consider *N* total miniprotein sequences, each with dissociation constants of *K*_*d,i*_, for *i* = 1, …*N*.

In a series of sequencing experiments, yeast cells displaying the designed miniproteins are incubated with a given concentration of fluorescently-labeled hemagglutinin (HA), and selected for sequencing using FACS. For a given HA concentration *L*, each sequence is read *M*_*i*_(*L*) times, for a total number of *M* (*L*) = Σ_*i*_ *M*_*i*_(*L*) sequence reads.

The initial pool of yeast cells display each binder with unknown efficiencies, but assume that the efficiencies are proportional to the number of sequences read in the control experiments performed in the absence of HA (the `rd0 seqMatch` numbers in the published data). Let *w*_*i*_ represent these baseline counts of sequence reads. We also assume that the fraction bound for each miniprotein is like that of a single-species binding curve :

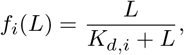

so that the probability of observing the set of *M*_*i*_ reads is given by the multinomial expression

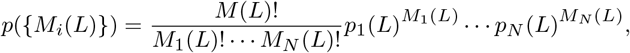

where *p*_*i*_(*L*) = *w*_*i*_*f*_*i*_(*L*)*/* Σ_*i*_ *w*_*i*_*f*_*i*_(*L*). The total probability of observing a set of sequence reads for experiments performed at *s* different HA concentrations *L* ∈ 𝒮 = [*L*_1_, *L*_2_, …, *L*_*s*_] is:

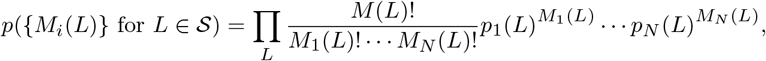

#### Bayesian inference of dissociation constants K_d,i_

The data *D* we are given for this inference problem are the sequencing read counts *D* = {*M*_*i*_(*L*)} for *L* ∈ [*L*_1_, *L*_2_, *L*_3_, *L*_4_] = [10^−6^ M, 10^−7^ M, 10^−8^ M, 10^−9^ M], the four HA concentrations used in the experiment. By Bayes’ theorem, the posterior probability of the set of dissociation constants is

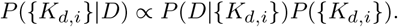

where *P* (*D*|{*K*_*d,i*_}) is the likelihood of observing the sequence read data *D* (the multinomial expression above), and *P* ({*K*_*d,i*_}) is a prior distribution. For the likelihood *P* (*D*|{*K*_*d,i*_}), we use the multinomial probability of observing the sequence reads described above. To model the prior distribution, we draw upon experimental ΔΔ*G* measurements of 1102 mutations in 57 protein-protein complexes published by Geng et al.[31] An analysis of this data shows that the distribution of Δ log_10_ *K*_*d*_ can be modeled as a Cauchy distribution (Figure S4):

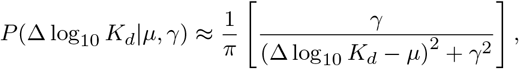

where *μ* is the mean and *γ* is the scale parameter (the half-width at half-maximum). The best-fit parameters are *μ* = ⟨Δ log_10_ *K*_*d*_⟩ = +0.532, and *γ* = 0.7045. Therefore, for the prior distribution of the set of {*K*_*d,i*_}, we use

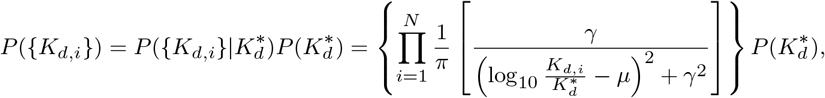

where 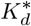 is the “baseline” *K*_*d*_ of wild-type miniprotein binder (A8, A13 or A18), and 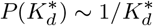 is a non-informative Jefferys’ prior.

To sample the posterior distribution of *P* ({*K*_*d,i*_}|*D*), MCMC was used in three stages. First, initial *K*_*d,i*_ values were set to 1.0 *μ*M, and 1M steps of MCMC were performed on a grid of log_10_ *K*_*d*_ values with spacing Δ log_10_ *K*_*d*_=0.3, and a move set where *n*_move_=5 randomly selected *K*_*d,i*_ values are chosen to move to adjacent grid positions (Figure S5a,b). Moves are accepted according to the Metropolis criterion, resulting in average acceptance probabilities of around *P*_acc_=0.34. Next a second round of MCMC is used to converge the *K*_*d,i*_ values near their most likely values. 1M MCMC steps were performed using a grid size of Δ log_10_ *K*_*d*_=0.1 and a *n*_move_=20 move set, resulting *P*_acc_ near 0.1, then decreasing to 0.05 as sampling progressed (Figure **??**c,d). Finally, a production run of 10M MCMC steps using a grid size of Δ log_10_ *K*_*d*_=0.05 and *n*_move_=20 move set was performed, resulting in *P*_acc_ ≈ 0.05 (Figure S5e). Whereas traces of the energy − ln *P* ({*K*_*d,i*_}|*D*) in the first two rounds show (mostly) monotonic decrease, fluctuations of the energy in the first round indicate equilibration (Figure S5f). Final estimates of the log_10_ *K*_*d*.*i*_ values and their uncertainties were calculated as the mean and standard deviation of the sampled values.

### 5.5 Sampling the distribution of sequences with multiple mutations

A simulated tempering algorithm was used to sample the probability distribution *P* (*s*) of sequences *s* with exactly *M* mutations, where *M* = 5 for A8, *M* = 7 for A13, and *M* = 6 for A18. We assume *P* (*s*) follows a Boltzmann distribution *P*_*β*_ (*s*) ∼ exp(−*βE*(*s*)),

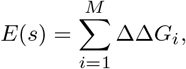

where the index *i* enumerates each mutation in the sequence, and *β* = 1*/RT*, where *R* is the gas constant and *T* = 298.15 K. To efficiently sample the entire sequence space, virtual replica exchange[63] is used to sample over six thermodynamic ensembles 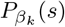, where *β*_*k*_ = *λ*_*k*_*β* for *λ* = [1, 0.8, 0.6, 0.4, 0.2, 0].

At each step in MCMC sampling protocol, a new sequence *s*^*′*^ is proposed with one of the *M* mutations randomly perturbed, and accepted with probability *P*_acc_ = min(1, exp(−*β*_*k*_ (*E*(*s*^*′*^) − *E*(*s*)). Every 10 steps, a virtual replica exchange from ensemble *k* to *l* = *k* ± 1 is additionally attempted, and accepted with probability *P*_acc_ = min(1, exp(−*β*_*k*_ (*E*(*s*^*′*^) − *E*(*s*)) exp(−(*β*_*l*_ − *β*_*k*_)*E*(*s*^*′*^)) exp(−(*β*_*k*_ − *β*_*l*_)*E*_*l*_)). The value of *E*_*l*_ is drawn from list of stored energies from ensemble *l*. The energies are stored every 10 MCMC steps, with a maximum number of 100K stored values (values are randomly replaced if this maximum is reached). Production-run MCMC was performed for 100M steps, with trajectory data saved every 100 steps.

## Data Availabilty

Molecular simulation input files and trajectory data are archived at https://osf.io/sj6mr/. Other data is available upon request.

## Code Availabilty

All code and analysis scripts are available at https://github.com/DJNovack/Massively Parallel Free Energy Calculations of HA Binders 2024/.

## Supplementary information

Supplementary Tables S1–S2, Supplementary Figures S1–S13.

## Acknowledgments

We thank the participants of Folding@home for making this work possible. We thank Christopher Bahl for many helpful discussions and for sharing with us the initial computational models of A8, A13, and A18 in complex with HA. This work is supported by the National Institutes of Health through NIH R01GM123296. Some calculations were performed on HPC resources supported in part by the National Science Foundation through NSF 1625061, the US Army Research Laboratory through W911NF-16-2-0189, NIH S10OD020095 and NIH R35GM132090.

## Supplementary Information

### A Supplementary Tables

**Table S1:** A8 miniprotein inter- and intraprotein contacts used in the salt-bridge and Laplacian centrality analysis.

| Residue | Contacts |
| --- | --- |
| T270 | Q140, S271, R272, T307, F308, D309 |
| S271 | Q140, R272, V273, V306, T307, F308 |
| R272 | V273, E305, V306, T307 |
| V273 | I143, D144, R274, A275, W304, E305, V306, F308 |
| R274 | N148, A275, Q303, W304, E305 |
| A275 | N148, N151, T276, Q303, W304 |
| T276 | E155, S277, K278, A281, G302, Q303, W304 |
| S277 | I154, K278, F279, A280, A281, W304 |
| K278 | F279, A280, A281, L282, W304 |
| F279 | L36, S63, L64, P65, V150, I154, A280, A281, L282, I283 |
| A280 | V150, N151, A281, L282, I283, A284 |
| A281 | L282, I283, A284, A285, V297, W304, V306 |
| L282 | I283, A284, A285, E286 |
| I283 | T90, V150, A284, A285, E286, I287 |
| A284 | T147, A285, E286, I287, A288, V306 |
| A285 | E286, I287, A288, R289, V295, V297, V306 |
| E286 | I287, A288, R289, E290 |
| I287 | I143, A288, R289, E290, F291 |
| A288 | R289, E290, F291, G292, Y293, V295, F308 |
| R289 | E290, F291, G292, Y293, V295 |
| E290 | H32, F291, G292 |
| F291 | I116, D117, G118, W119, T139, G292, Y293 |
| G292 | Y293, T294 |
| Y293 | Q136, T294, V295, F308, D309 |
| T294 | V295, D296, T307, F308, D309 |
| V295 | D296, V297, V306, T307, F308 |
| D296 | V297, Q298, E305, V306, T307 |
| V297 | Q298, W304, E305, V306 |
| Q298 | E299, V300, Q303, W304, E305 |
| E299 | V300, N301, G302, Q303, W304 |
| V300 | N301, G302, Q303, E305 |
| N301 | G302, Q303 |
| G302 | Q303, W304 |
| Q303 | W304, E305 |
| W304 | E305, V306 |
| E305 | V306, T307 |
| V306 | T307, F308 |
| T307 | F308 |
| F308 | I143, D309 |
| D309 |  |

**Table S2:** A13 miniprotein inter- and intraprotein contacts used in the salt-bridge and Laplacian centrality analysis.

| Residue | Contacts |
| --- | --- |
| C270 | I271, E272, H307, V308, C309 |
| I271 | E272, I273, L306, H307, V308 |
| E272 | I273, S274, R305, L306 |
| I273 | S274, V275, W288, C304, R305, L306, V308 |
| S274 | V275, N303, C304 |
| V275 | N148, T276, T277, C281, G302, N303, C304, L306 |
| T276 | E155, T277, P278, A280, C281, G302, N303, C304 |
| T277 | I154, P278, F279, A280, C281, G302 |
| P278 | F279, A280, C281, Q282, D299, G302 |
| F279 | L36, S63, L64, P65, V150, I154, A280, C281, Q282, I283 |
| A280 | V150, N151, C281, Q282, I283, A284 |
| C281 | Q282, I283, A284, A285, I297, C304, L306 |
| Q282 | I283, A284, A285, E286, I297 |
| I283 | T90, V150, A284, A285, E286, I287 |
| A284 | T147, A285, E286, I287, W288, L306 |
| A285 | E286, I287, W288, R289, V295, I297 |
| E286 | I287, W288, R289, A290 |
| I287 | I143, W288, R289, A290, F291 |
| W288 | I143, R289, A290, F291, G292, Y293, V295, V308 |
| R289 | A290, F291, G292, Y293 |
| A290 | I116, F291, G292 |
| F291 | I116, D117, G118, W119, T139, I143, G292, Y293 |
| G292 | Y293, E294 |
| Y293 | Q136, E294, V295, V308, C309 |
| E294 | V295, K296, H307, V308, C309 |
| V295 | K296, I297, L306, H307, V308 |
| K296 | I297, D298, R305, L306, H307 |
| I297 | D298, D299, C304, R305, L306 |
| D298 | D299, D300, N303, C304, R305, H307 |
| D299 | D300, N301, G302, N303, C304 |
| D300 | N301, G302, N303, C304, R305 |
| N301 | G302, N303 |
| G302 | N303, C304 |
| N303 | C304, R305 |
| C304 | R305, L306 |
| R305 | L306, H307 |
| L306 | H307, V308 |
| H307 | V308, C309 |
| V308 | C309 |
| C309 |  |

**Table S3:** A18 miniprotein inter- and intraprotein contacts used in the salt-bridge and Laplacian centrality analysis.

| Residue | Contacts |
| --- | --- |
| C270 | Q271, D272, R307, C308, C309 |
| Q271 | D272, Y273, V306, R307, C308 |
| D272 | Y273, T274, V306, R307 |
| Y273 | T274, F275, L288, C304, R305, V306, C308 |
| T274 | F275, C304 |
| F275 | D144, T147, T276, D277, C281, A284, G302, E303, C304, V306 |
| T276 | N151, D277, P278, A280, C281, G302 |
| D277 | I154, P278, F279, A280, C281, G302, C304 |
| P278 | F279, A280, C281, Q282, T299, G302, C304 |
| F279 | L36, S63, L64, P65, V150, I154, A280, C281, Q282, I283 |
| A280 | V150, N151, C281, Q282, I283, A284 |
| C281 | Q282, I283, A284, A285, V297, C304, V306 |
| Q282 | I283, A284, A285, E286, R289, V297 |
| I283 | T90, V150, A284, A285, E286, I287 |
| A284 | T147, A285, E286, I287, L288 |
| A285 | E286, I287, L288, R289, C295, V297, V306 |
| E286 | I287, L288, R289, D290 |
| I287 | W119, I143, L288, R289, D290, F291 |
| L288 | R289, D290, F291, G292, Y293, C295, C308 |
| R289 | D290, F291, G292, Y293, C295, T296 |
| D290 | I116, F291, G292 |
| F291 | I116, D117, G118, W119, T139, I143, G292, Y293 |
| G292 | Y293, D294 |
| Y293 | Q136, D294, C295, C308, C309 |
| D294 | C295, T296, R307, C308, C309 |
| C295 | T296, V306, R307, C308 |
| T296 | V297, Q298, R305, V306, R307 |
| V297 | Q298, T299, R305, V306 |
| Q298 | T299, N300, E303, C304, R305 |
| T299 | N300, N301, E303, C304 |
| N300 | N301, G302, E303 |
| N301 | G302, E303 |
| G302 | E303, C304 |
| E303 | C304, R305 |
| C304 | R305, V306 |
| R305 | V306, R307 |
| V306 | R307, C308 |
| R307 | C308, C309 |
| C308 | C309 |
| C309 |  |

### B Supplementary Figures

**Fig. S1:**
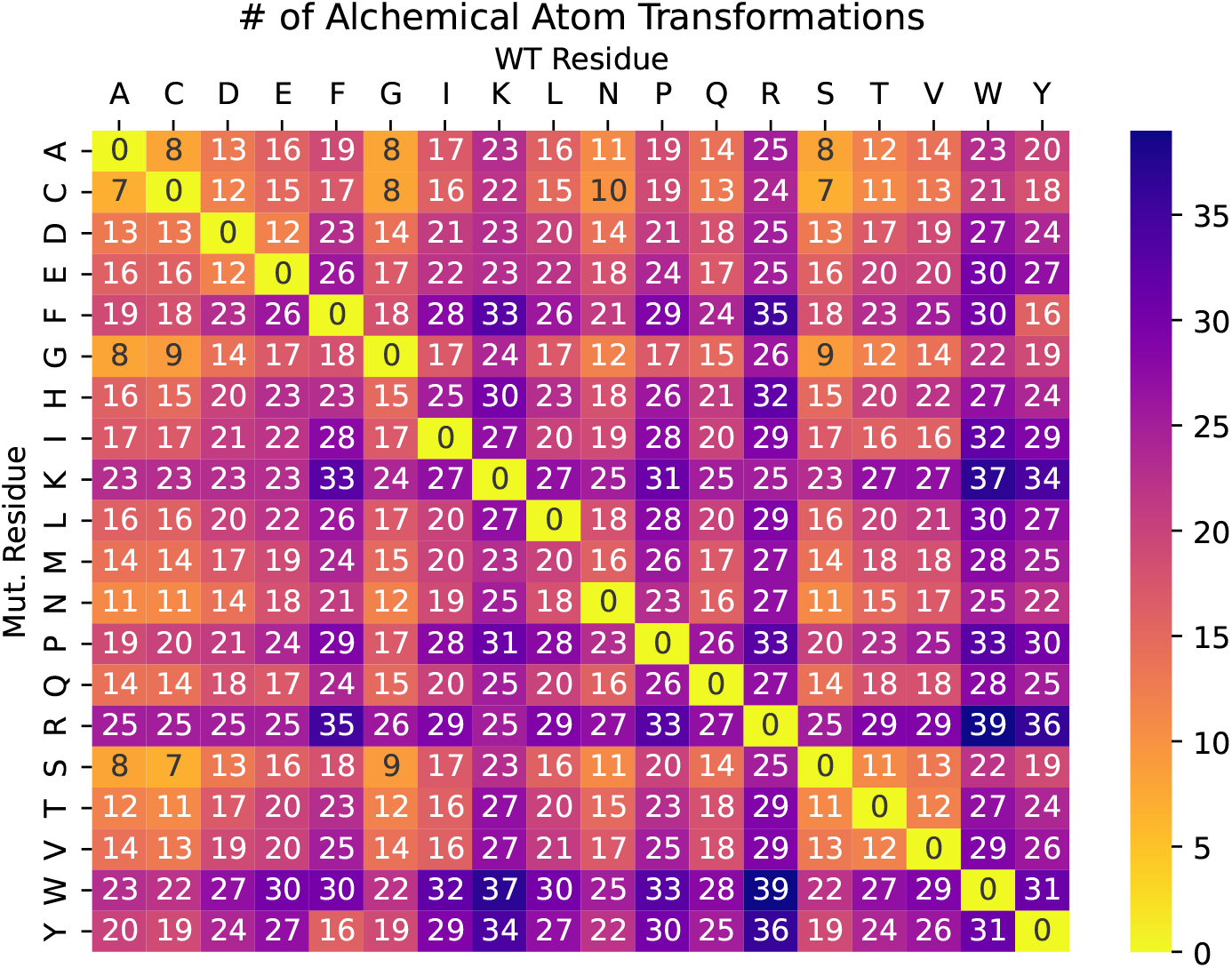
Number of atoms involved in alchemical transformations, arranged by wild-type and mutant amino acid identity. Atom counts include hydrogens.

**Fig. S2:**
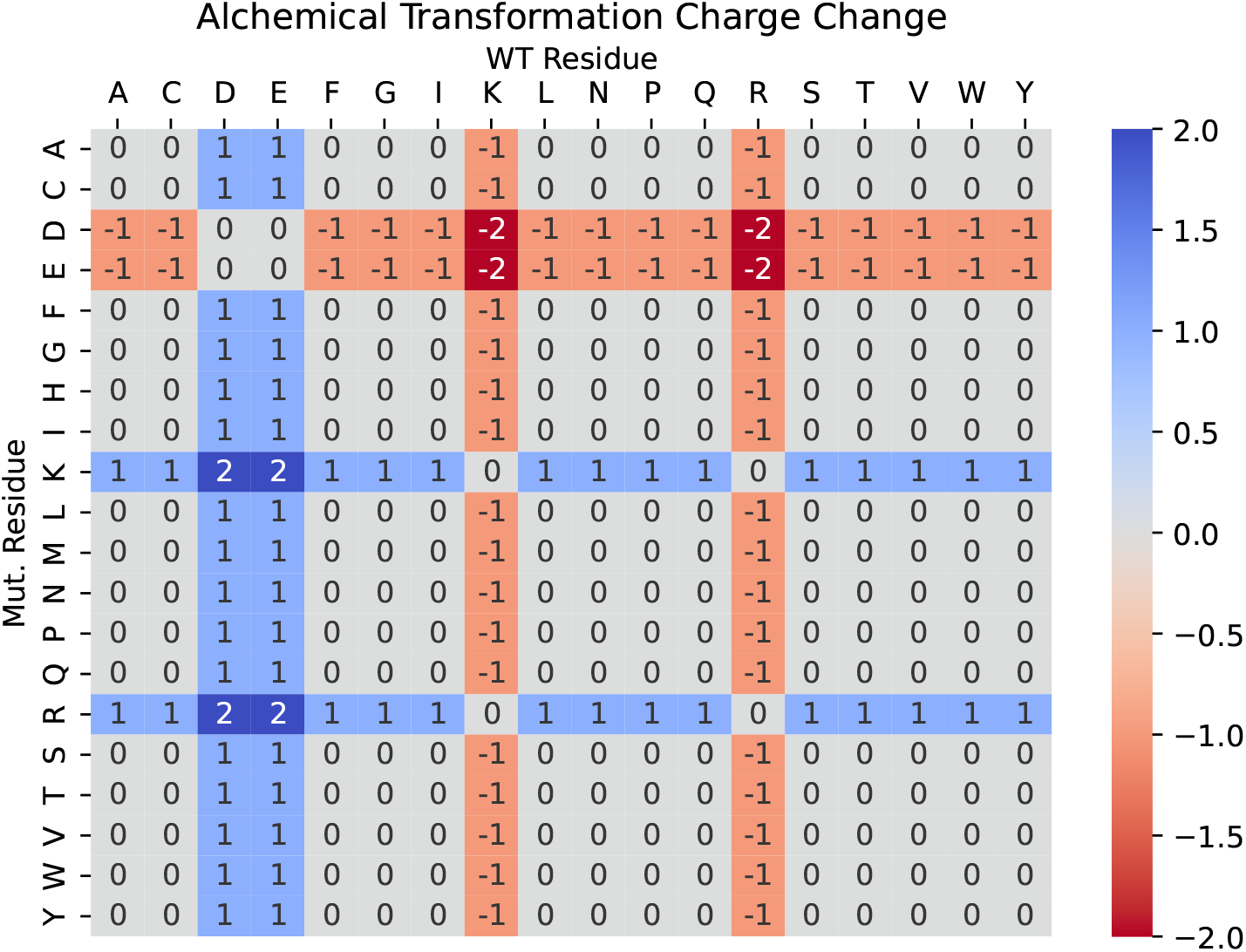
Net charge changes involved in alchemical transformations, arranged by wild-type and mutant amino acid identity.

**Fig. S3:**
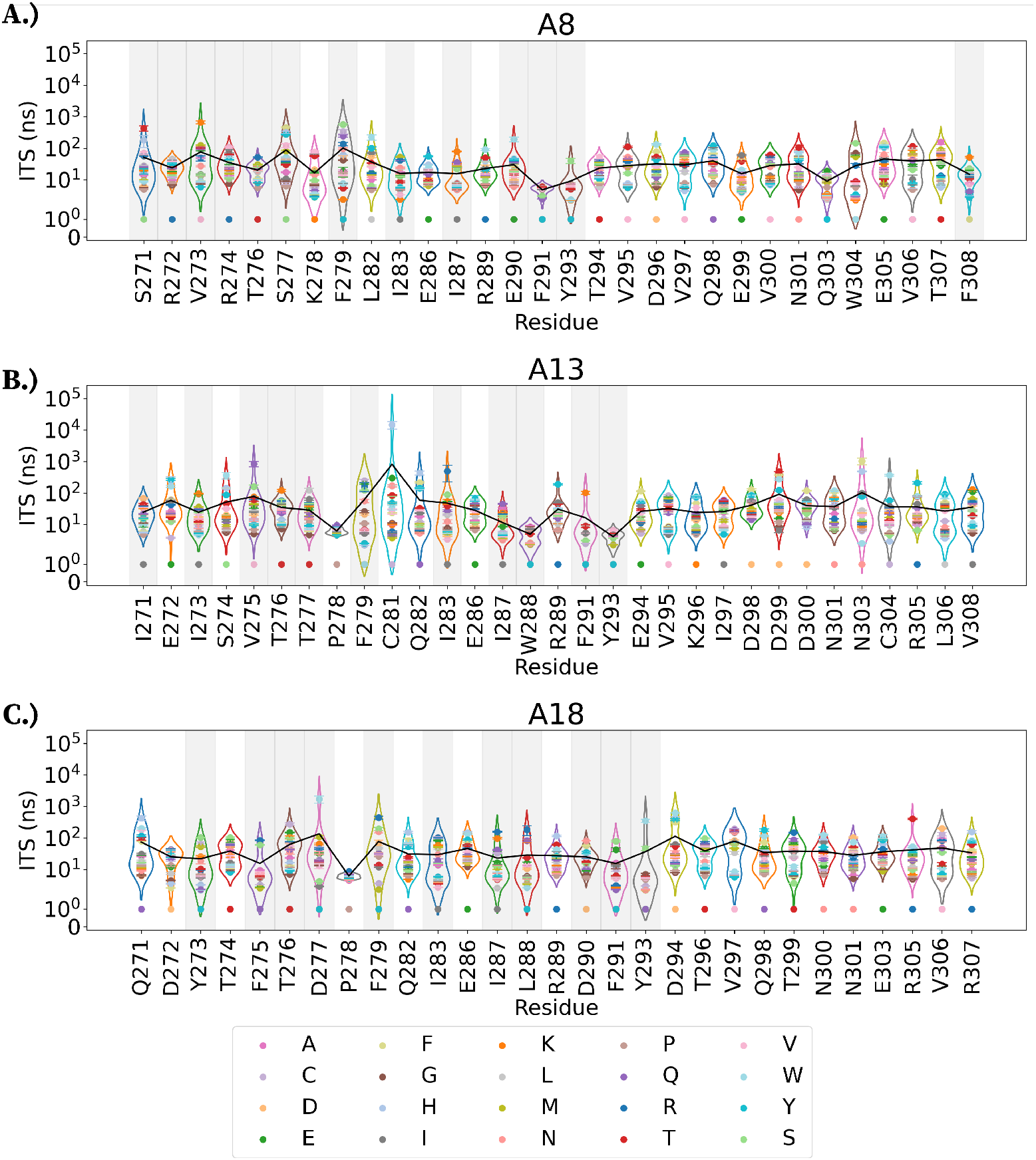
Average implied timescales of side-chain dynamics at each residue position. Violin plots of implied timescales for mutations to all other amino acids at each residue position for miniproteins (A) A8, (B) A13, and (C) A18. Error bars represent the standard error of the mean. Gray shaded regions are interfacial residues. Mean implied timescales of each residue is shown as a solid black line. Markers at the bottom of each violin denote the wild-type residue identity.

**Fig. S4:**
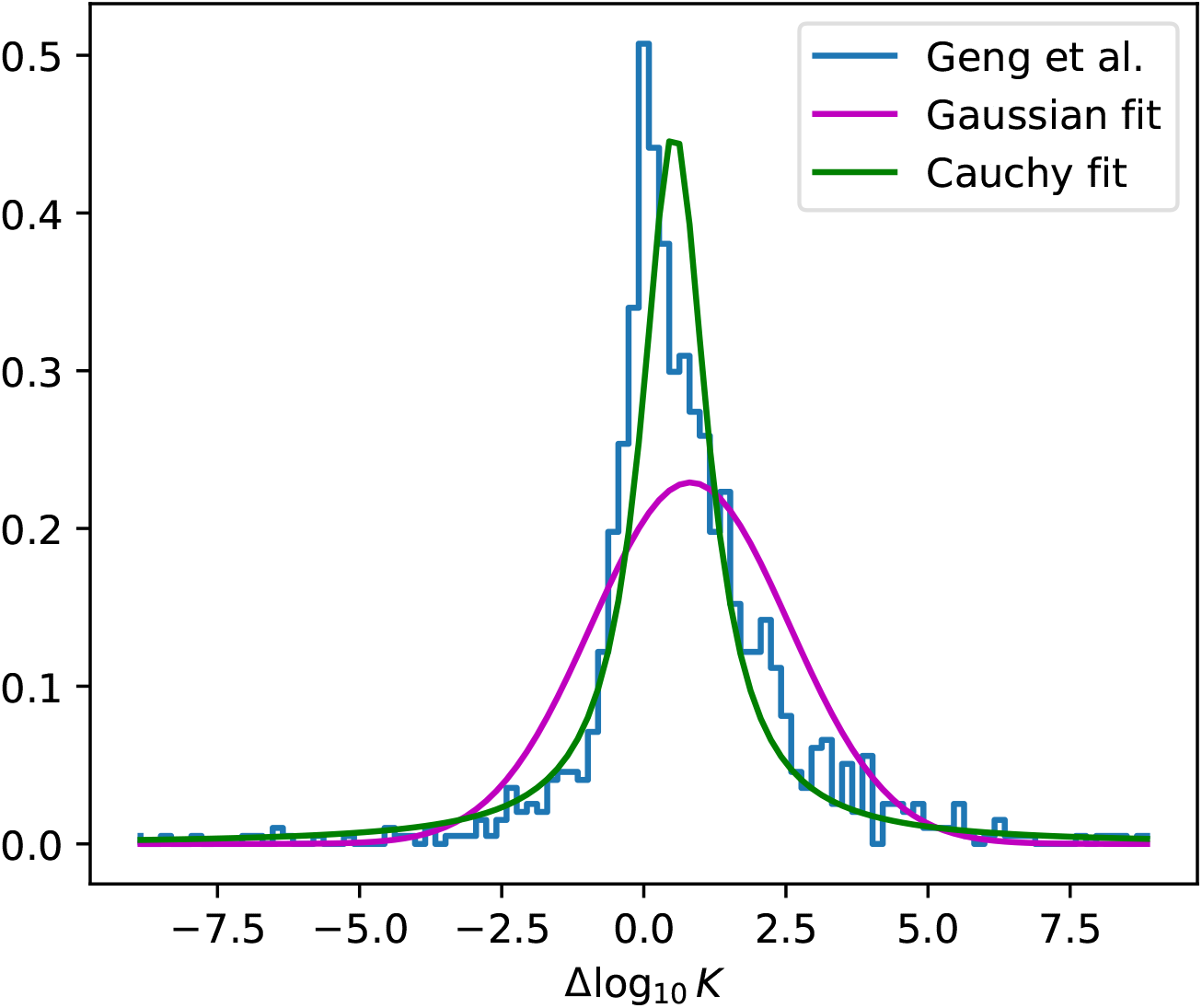
The experimental distribution of Δ log_10_ *K*_*d*_ for 1102 single point mutations in 57 protein complexes from SKEMPI 1.1 database by Geng et al. 2017 (blue), super-imposed with best-fit Gaussian distribution (magenta, *μ*=0.819, *σ*=1.741) and Cauchy distribution (green, *μ*=0.532, *γ*=0.7045). The Cauchy distribution is better able to capture the tails of the experimental distribution.

**Fig. S5:**
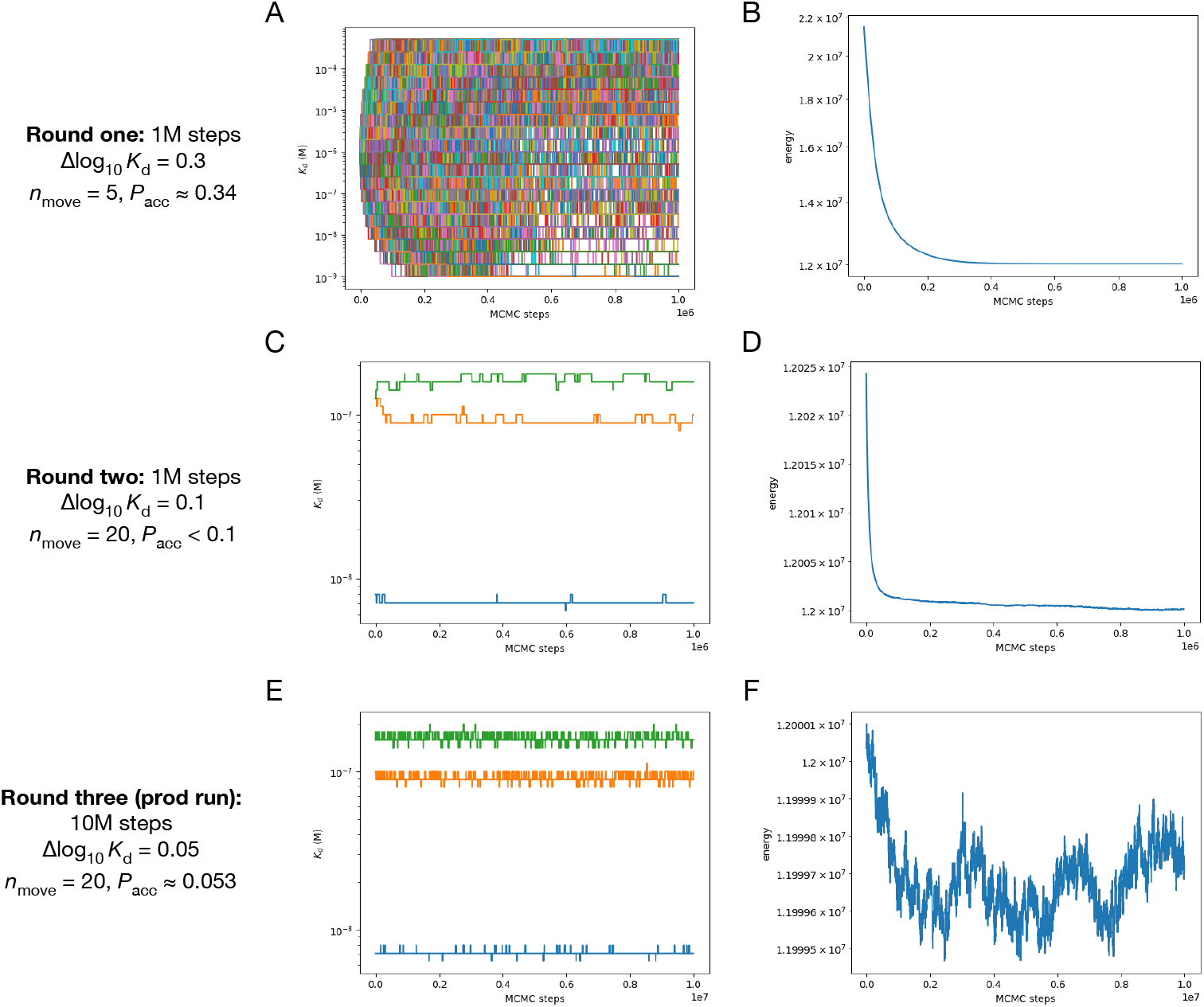
MCMC sampling of the Bayesian posterior *P* ({*K*_*d,i*_}|*D*). (a) Traces of all *K*_*d,i*_ versus MCMC step in the first round of sampling. (b) Traces of the energy −ln *P* ({*K*_*d,i*_} |*D*) in the first round of sampling. (c) Traces of three example *K*_*d,i*_ values (for A8, A13 and A18 sequences) versus MCMC step in the second round of sampling. (d) Energy trace in the second round of sampling. (e) Traces of three example *K*_*d,i*_ values (for A8, A13 and A18 sequences) versus MCMC step in the final production-run round of sampling. (f) In this final round of sampling, energy traces fluctuate at equilibrium.

**Fig. S6:**
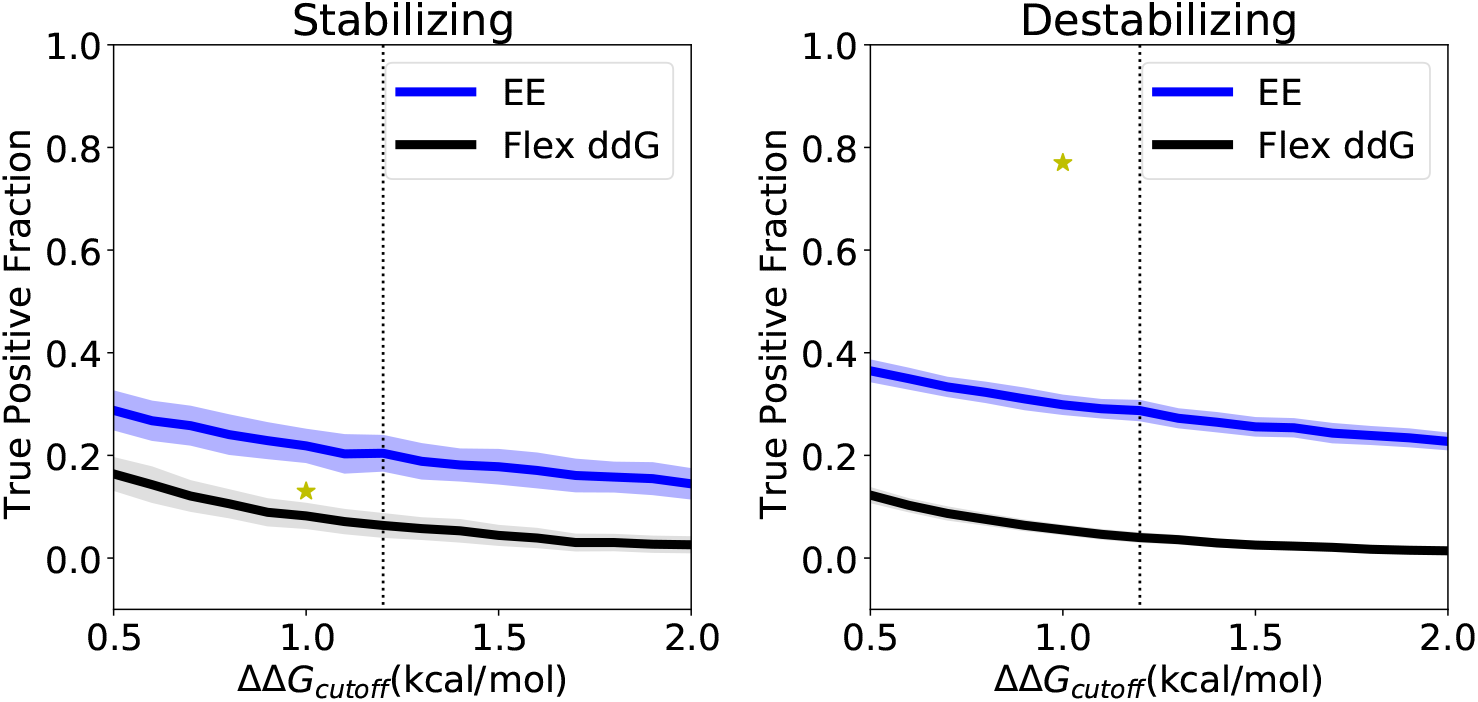
A recent study by Cao et al. (2022) measured experimental ΔΔ*G* for 12 *de novo* designed miniprotein binder SSMs and compared against predictions from Rosetta using a classification threshold of 1.0 kcal/mol.[3] True positive rates of stabilizing (0.13, left) and destabilizing mutations (0.77, right) from Cao et al. (2022) are show as gold stars, superimposed on Figures 8f and 8g from the main text.

**Fig. S7:**
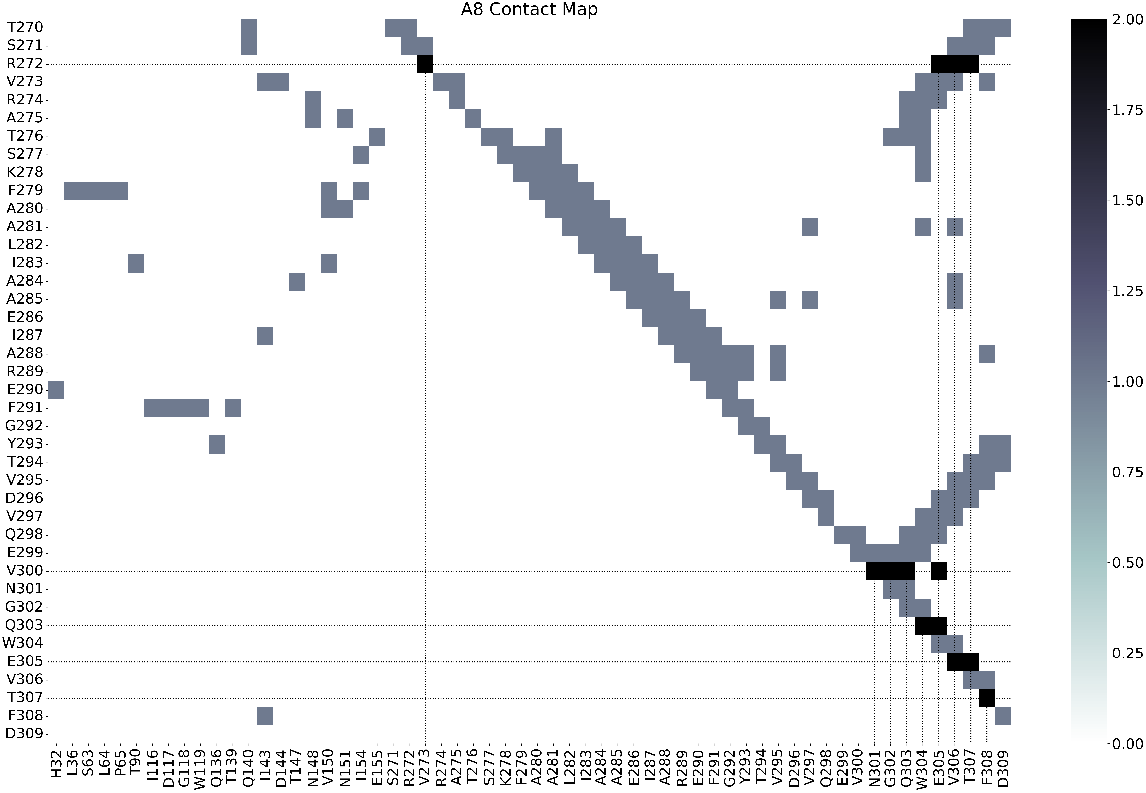
A8 Contact Map. Pairwise residue contact map between A8 miniprotein (vertical axis) and HA2 and itself (horizontal axis). The vertical lines mark residues that make contact with miniprotein residues observed to be mutated in the affinity-matured variant (horizontal lines). Grey cells denote contacts, while black cells denote contacts at positions mutated in the affinity-matured variant.

**Fig. S8:**
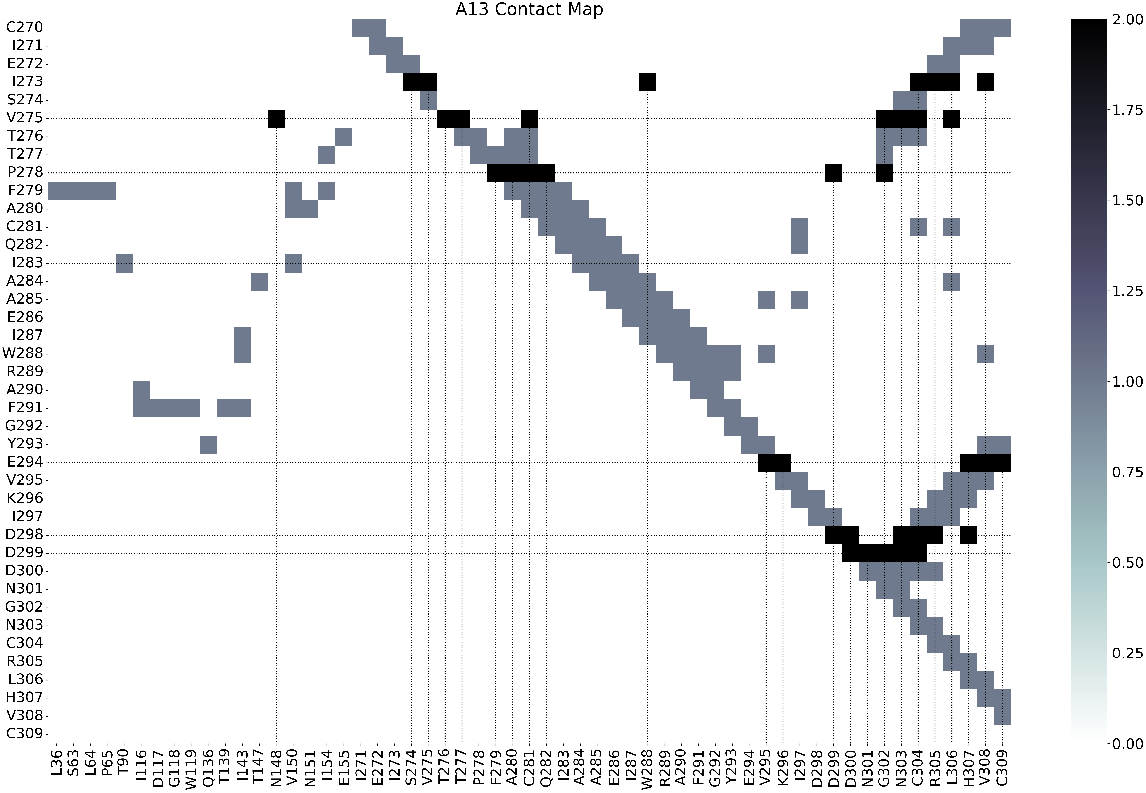
A13 Contact Map. Pairwise residue contact map between A13 miniprotein (vertical axis) and HA2 and itself (horizontal axis). The vertical lines mark residues that make contact with miniprotein residues observed to be mutated in the affinity-matured variant (horizontal lines). Grey cells denote contacts, while black cells denote contacts at positions mutated in the affinity-matured variant.

**Fig. S9:**
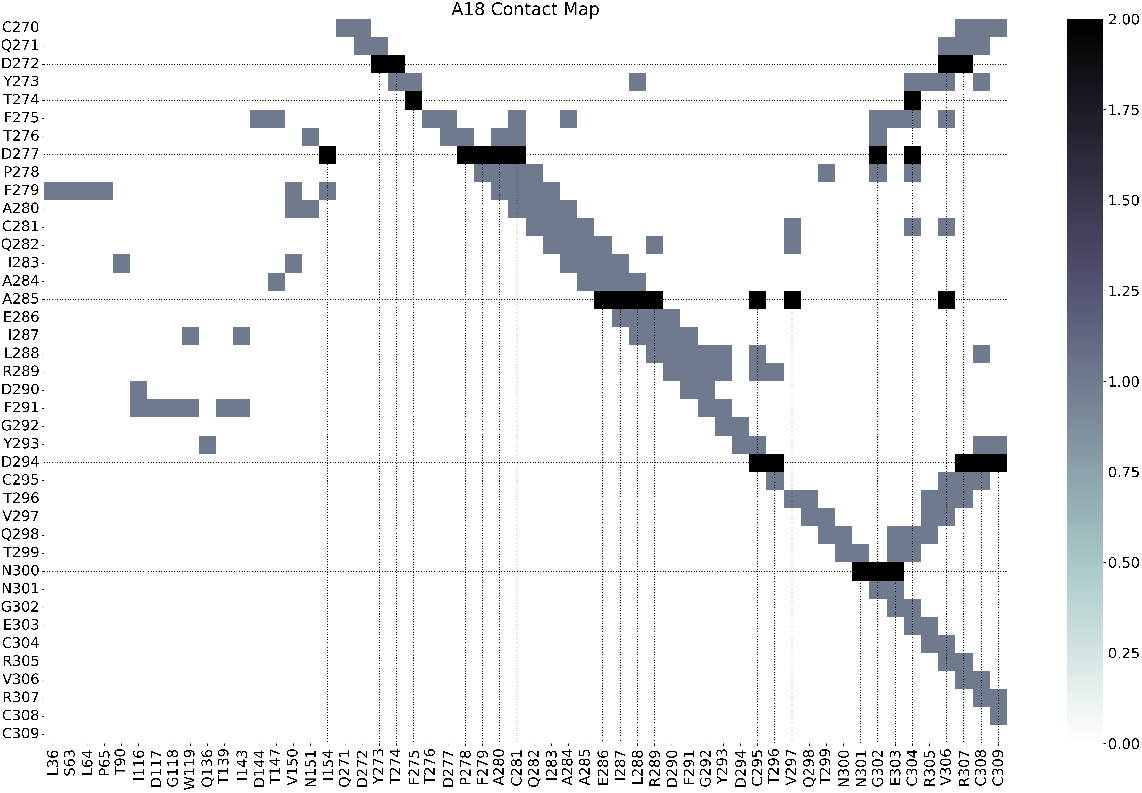
A18 Contact Map. Pairwise residue contact map between A18 miniprotein (vertical axis) and HA2 and itself (horizontal axis). The vertical lines mark residues that make contact with miniprotein residues observed to be mutated in the affinity-matured variant (horizontal lines). Grey cells denote contacts, while black cells denote contacts at positions mutated in the affinity-matured variant.

**Fig. S10:**
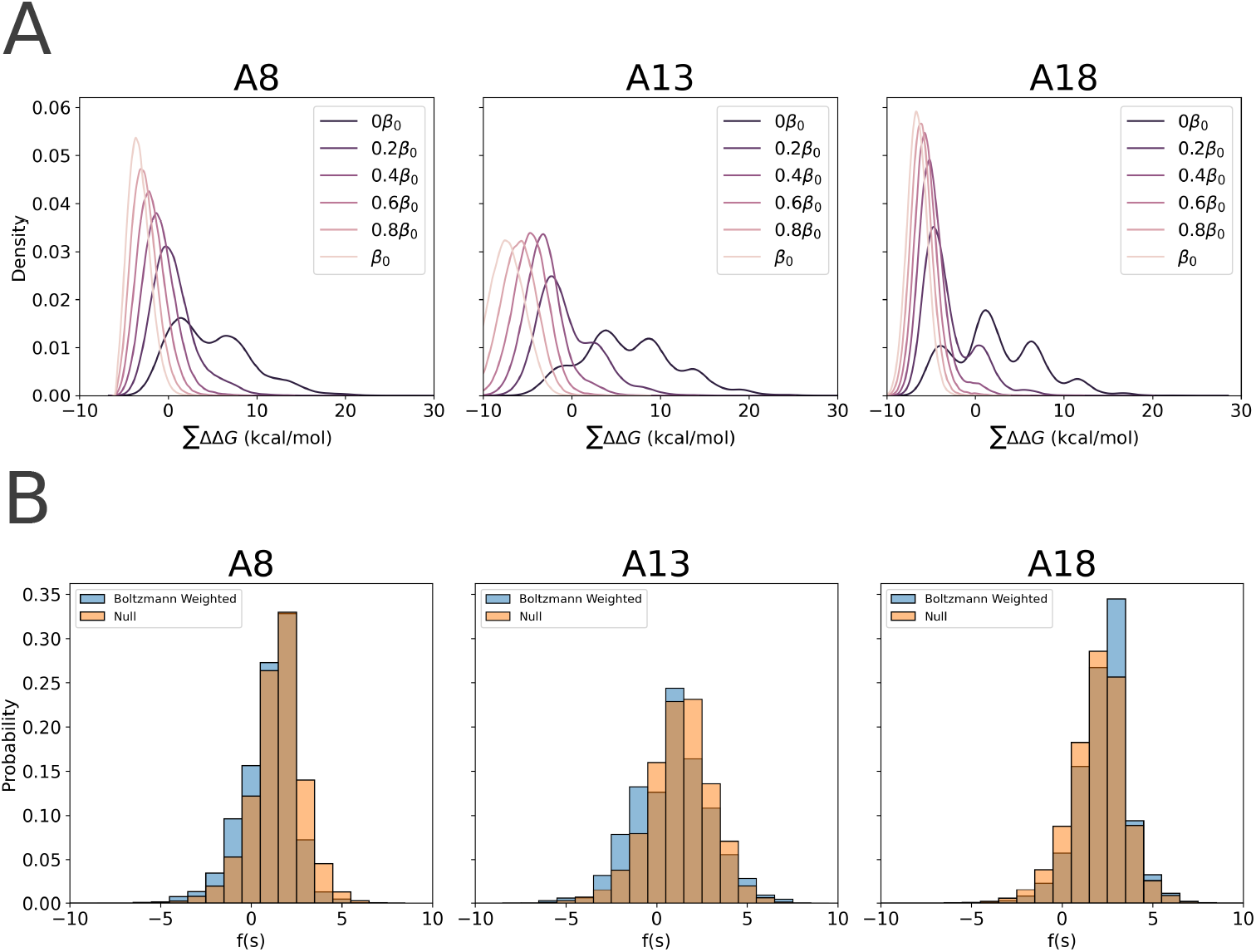
Simulated Tempering MCMC to Investigate Salt Bridge Effects. (A) Density of states of all 5-(A8) and 6-residue (A13 and A18) mutant sequences from thermodynamic ensembles at various inverse temperatures *β*_*k*_ = *λ*_*k*_*β*, where *λ* = [1, 0.8, 0.6, 0.4, 0.2, 0], sampled using simulated tempering with virtual replica exchange. The reference inverse temperature was *β* = 1*/RT*, where *T* = 300 K. (B) Histogram of sampled values of *f* (*s*) quantifying net electrostatic effects under (blue) the Boltzmann distribution at inverse temperature *β*, and (orange) the null distribution at zero inverse temperature.

**Fig. S11:**
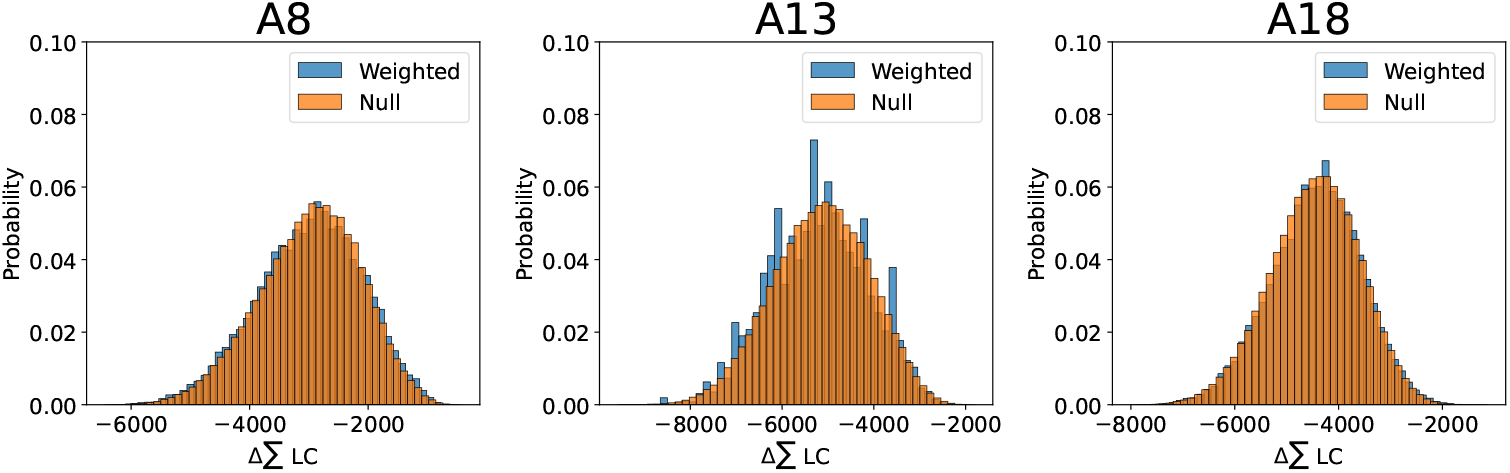
Histograms of Laplacian Centrality Differences for all 5-(A8) and 6-residue (A13 and A18) mutant sequences, sampled from (blue) the Boltzmann distribution at inverse temperature *β*, and (orange) the null distribution at zero inverse temperature.

**Fig. S12:**
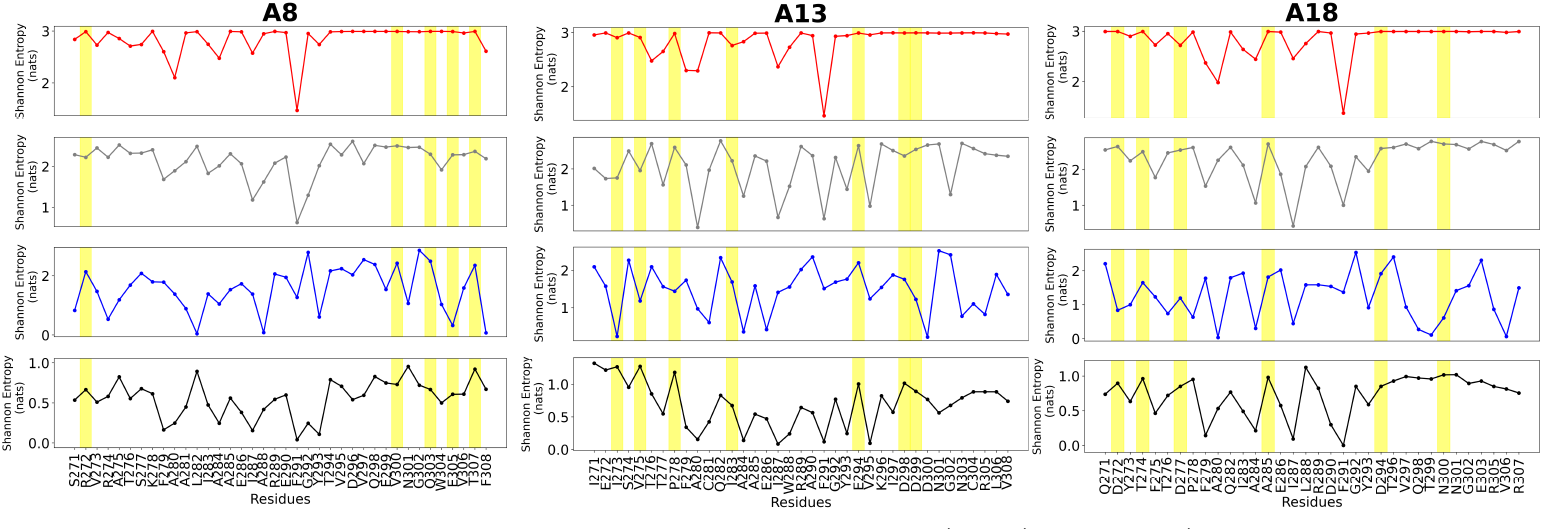
Shannon entropy profiles of miniproteins A8, A13 and A18 computed using ΔΔ*G* estimates from Flex ddG (red, top row), inferred experimental values (gray, second row), expanded ensemble estimates (blue, third row), and frequencies from the simulated tempering sampling (black, bottom row).

